# Similar but different: ERP evidence on the processing of mental and physical experiencer verbs in Malayalam

**DOI:** 10.1101/2025.03.12.642939

**Authors:** S. Shalu, R. Muralikrishnan, Anna Merin Mathew, Kamal Kumar Choudhary

**Author notes:** Correspondence: R. Muralikrishnan, Max Planck Institute for Empirical Aesthetics, Grüneburgweg 14, 60322 Frankfurt am Main, Germany.

## Abstract

The study reported here was conducted to investigate the neurophysiological correlates of processing two types of subject experiencer verbs, namely mental experiencer verbs and physical experiencer verbs in Malayalam, a South Dravidian language. Event-related brain potentials (ERPs) were recorded as twenty-eight first-language speakers of Malayalam read intransitive sentences with the two types of experiencer verbs. Critical stimuli were either fully acceptable, whereby the subject case matched the requirements of the verb, or unacceptable, whereby the subject case violated the requirements of the verb. A linear mixed-models analysis confirmed negativity effects in the time window 400–800 ms for mental and physical experiencer verbs. Post-hoc analyses revealed that the negativity peaked relatively early for mental experiencer verbs, whereas relatively late for physical experiencer verbs. Further, the sentence-final acceptability of trials modulated the ERPs in non-anomalous conditions but not in violation conditions, and this modulation qualitatively differed between mental and physical experiencer verbs. These results suggest that, whilst a qualitatively similar mechanism is involved in the processing of both kinds of experiencer verbs, subtle but robust differences are inherent in processing mental versus physical experiencer verbs in Malayalam.

## 1. Introduction

Experiencers are special, both cognitively and linguistically (Landau, 2009). Understanding and categorizing entities and activities as experiencers or non-experiencers are fundamental to how we navigate and interpret the world. This cognitive significance of experiencers has some consequences for the grammar of a language (Dabrowska,1994; Jayaseelan, 2004; Landau, 2009; Shimoyoshi, 2015). Experiencer predicates are used across languages to convey feelings, whether they pertain to emotions such as happiness, sadness, anger, etc. (Levin & Grafmiller, 2013), or bodily sensations like pain, hunger, cold, etc. (Fleischhauer, 2016). A key feature of these verbs is that one of the participants is an experiencer (Dowty, 1991), who can be either a physical experiencer or a mental experiencer. Depending on the realization of this experiencer participant, experiencer verbs are broadly categorized into two classes: Subject Experiencer verbs (SE) and Object Experiencer verbs (OE) (Postal, 1971; Belletti & Rizzi, 1988; Lakoff, 1971; Jackendoff, 1972; Levin, 1993; M.A. Anne Temme, 2018). Such verbs are found in various languages (1-4).

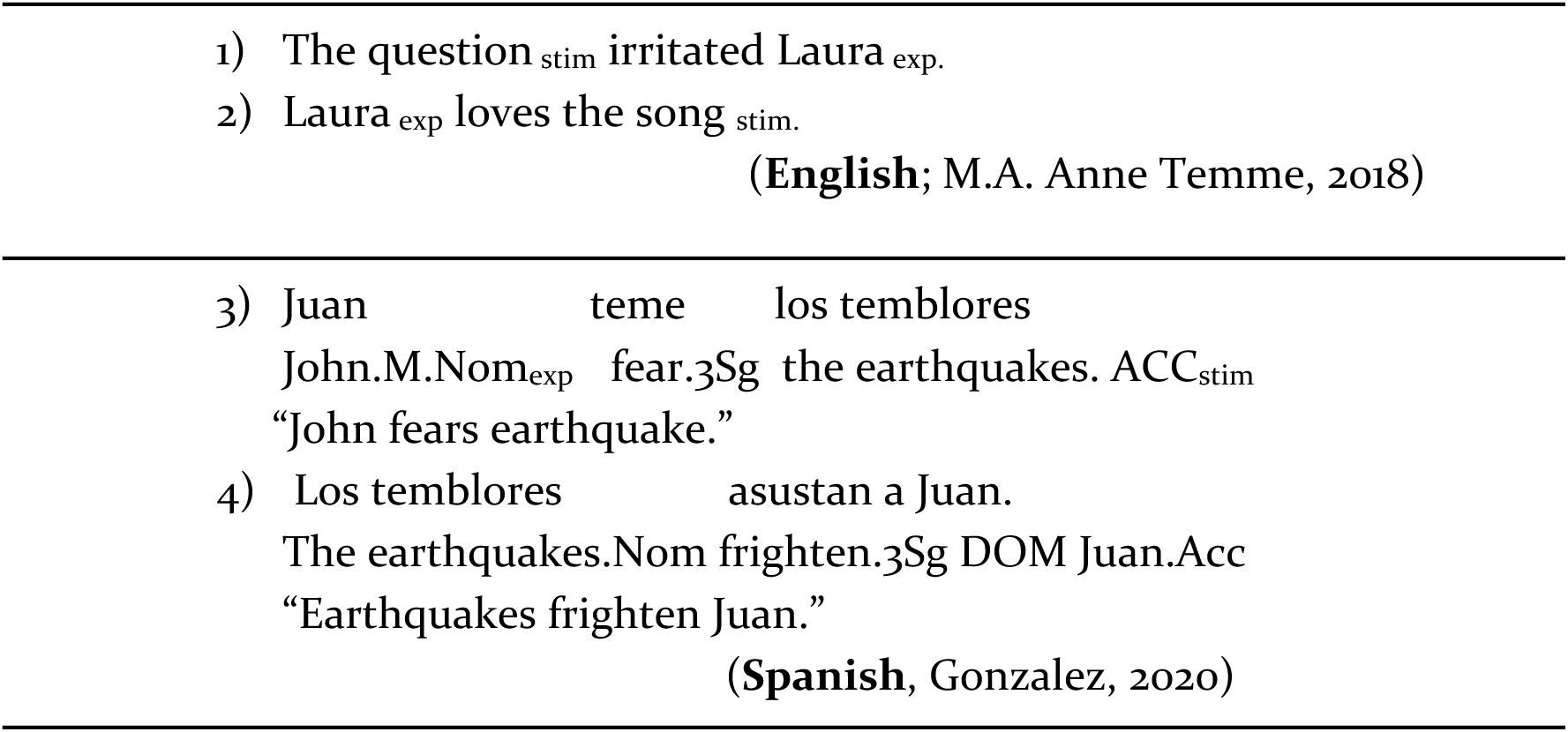

These verbs differ from other verb classes because of their internal (semantic) structure, characterized by being static rather than event-oriented (Davidson, 1967; Dowty, 1979; Jackendoff, 1991; Rappaport Hovav and Levin, 2015), their argument structure and realization, which encode the verb’s obligatory argument and assignment of thematic roles to the syntactic constituents (Postal, 1971; Belletti & Rizzi, 1988; Lakoff, 1970; Jackendoff, 1990; Levin, 1993).

Experiencer verbs have been an important topic of research in linguistic theories and language processing studies due to their unique syntactic (Belletti, & Rizzi, 1988; Postal, 1971) and semantic (Grimshaw, 1990; Dowty, 1991; Croft,1993; Pesetsky,1995; Van Valin, 2005) features. Previous processing research on experiencer verbs (or psych-verbs) have broadly addressed questions about argument structure and realization, semantic complexity, and embodied cognition (for an overview, see Brennan, 2015).

Several behavioural studies have investigated the processing differences between object experiencer verbs and subject experiencer verbs in individuals with language impairments (Grodzinsky, 1995; Bretta and Campbell, 2001; Thompson and Lee, 2009, etc.) as well as in those without such impairments (Cupple, 2002; Lamers, 2007; Kretzschmar et al.2012; Gattei et al. 2022; Mateu et al. 2022; Wilson and Dillon, 2022, etc). The results of these studies showed that subject-experiencer verbs are easier to process than object-experiencer verbs, as object experiencer verbs show longer reaction time, higher error rates and significantly more fixations and regression during reading. This is because of the less complicated alignment encoding thematic role and grammatical function (Cupple, 2002; Kretzschmar et al., 2012; Gattei et al., 2015; Gattei et al., 2018; Gattei et al., 2022; Mateu, 2022; Wilson & Dillon, 2022; ) and low semantic complexities associated with subject experiencer verbs (Mckoon and Macfarland, 2002; Gennari and Poeppel, 2003; Frisson and Frazier, 2005; Brennan and Pylkkänen, 2008; 2010). However, studies on the subcategories of object experiencer verbs have shown that the processing of object experiencer verbs varies based on how their arguments are expressed, such as in accusative versus dative constructions. Dative object experiencer verbs generally show higher processing costs compared to accusative ones due to differences in linking, i.e., direct vs. inverse syntax to semantic linking (Gattei et al., 2022).

Experiencer verbs have been employed in research on embodied cognition to examine how sensory-motor information contributes to semantic representation. Specifically, several neurophysiological studies have investigated the activation of motor cortices during the comprehension of concrete verbs (action verbs) and abstract verbs (experiencer verbs) (Tettamanti et al., 2005; Bedney et al., 2008; 2012; Javier Rodriguez-Ferreiro, 2011; Kemmerer et al., 2013; Murakai et al., 2020, etc.). In these studies, experiencer verbs were either used as a control condition due to their lack of concrete motor and sensory associations, or they were compared to other types of abstract verbs. Findings from some of these studies indicate that various types of abstract verbs are processed differently depending on whether they pertain to emotional states, mental processes, or non-bodily states (Muraki et al., 2020). These differences are said to arise from the different representational systems (sensorimotor vs. linguistic) by each verb type.

Several ERP studies have employed experiencer verbs in investigations related to reanalysis, primarily because of the non-canonical thematic ordering inherent to such verbs and the word order preferences associated with them. These studies have mainly investigated object experiencer verbs because of their distinct reverse-linking nature and semantic complexities, with only a few studies examining the processing of subject experiencer verbs.

The studies on object experiencer verbs, many of which are on German, capitalised on the fact that dative object experiencer verbs in German are said to project a dative initial base order because the dative marked experiencer is argued to outrank the nominative case marked theme (Wunderlich, 1997; 2003; Grimshaw,1990). Bornkessel et al. (2002) and Bornkessel et al. (2003) utilized this aspect of German experiencer verbs to investigate the processing differences between object experiencer verbs and action verbs. Clause-final object experiencer verbs engendered a late positivity as opposed to active verbs, regardless of case ambiguity or word order, which was interpreted as resulting from a reanalysis occurring when the preferred thematic ordering (first argument as thematically highest) is contradicted by the verb. Similar reanalysis related late positivities have been reported in Italian (Dröge et al., 2014) and Spanish (Gattei et al., 2015) for the processing of object experiencer verbs. Bornkessel et al. (2004) compared object experiencer verbs with action verbs to investigate whether the lexical information associated with object experiencer verbs modulates the grammatical function reanalysis. The study revealed that clause-final object experiencer verbs engendered less pronounced negativity compared to action verbs, suggesting that their lexical information facilitates reanalysis. Additionally, a LAN was observed for object experiencer verbs in nominative initial structures compared to dative initial ones, which was said to reflect the mismatch between thematic hierarchy (Experiencer>Theme) and case hierarchy (Nominative>Dative). However, in ERP studies in which the verb is not clause-final but instead precedes the disambiguating argument, object-experiencer verbs align more closely with action verbs (Schlesewsky & Bornkessel, 2006).

Interest in the processing patterns of subject-experiencer verbs emerged after the proposition by Bornkessel-Schlesewsky et al. (2011) that different verb types within a language might evoke distinct ERP responses to thematic reversal anomalies (henceforth TRA). This prompted studies that focused on subject experiencer verbs in the context of TRA (Bourguignon et al., 2012; Kyriaki et al., 2020). Findings from these studies suggest that subject experiencer verbs differ qualitatively from other verb types, particularly action verbs, due to differences in their thematic and aspectual structure.

The overall findings from studies on experiencer verbs can be summarised thus: a) subject experiencer verbs are easier to process than object experiencer verbs; b) object experiencer verbs with differences in how the experiencer argument is realized show concomitant processing differences, whereby dative experiencers incur higher processing costs than accusative experiencers.

Subject experiencer verbs express different kinds of experiences including physical experiences such as hunger, thirst, cold, etc., as in sentence (5), and mental experiences such as happiness, sadness, etc., as in sentence (6). However, it is not clear whether and how the processing of various types of subject experiencer verbs differs. This is because there is hardly any study to date that investigated the processing of various types of subject experiencer verbs, particularly experiencer verbs indicating different kinds of experiences.

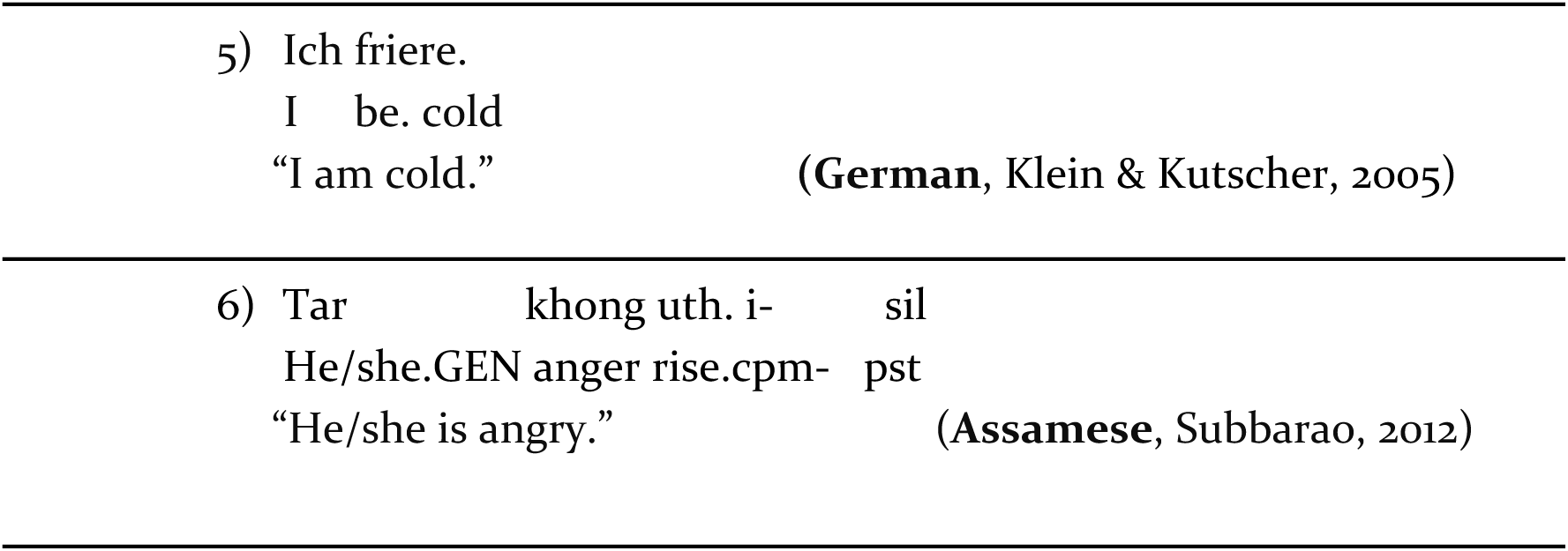

|  |  |
| --- | --- |
| 5) Ich friere. |  |
| I be. cold |  |
| “I am cold.” | (German, Klein & Kutscher, 2005) |

|  |  |  |  |
| --- | --- | --- | --- |
| 6) Tar | khong uth. i- | sil |  |
| He/she.GEN | anger rise.cpm- | pst |  |
| “He/she is angry.” |  |  | (Assamese, Subbarao, 2012) |

Whilst such verbs are present in several languages of the world, the case of Malayalam, a south Dravidian language with a canonical SOV word order, is particularly interesting in this regard because it clearly distinguishes between verbs denoting a mental experience (henceforth mental experiencer verbs) and those denoting a physical experience (henceforth physical experiencer verbs) in its syntax (Jayaseelan, 2004). Malayalam employs different case markings on the experiencer arguments based on the type of experience indicated by the experiencer verbs.

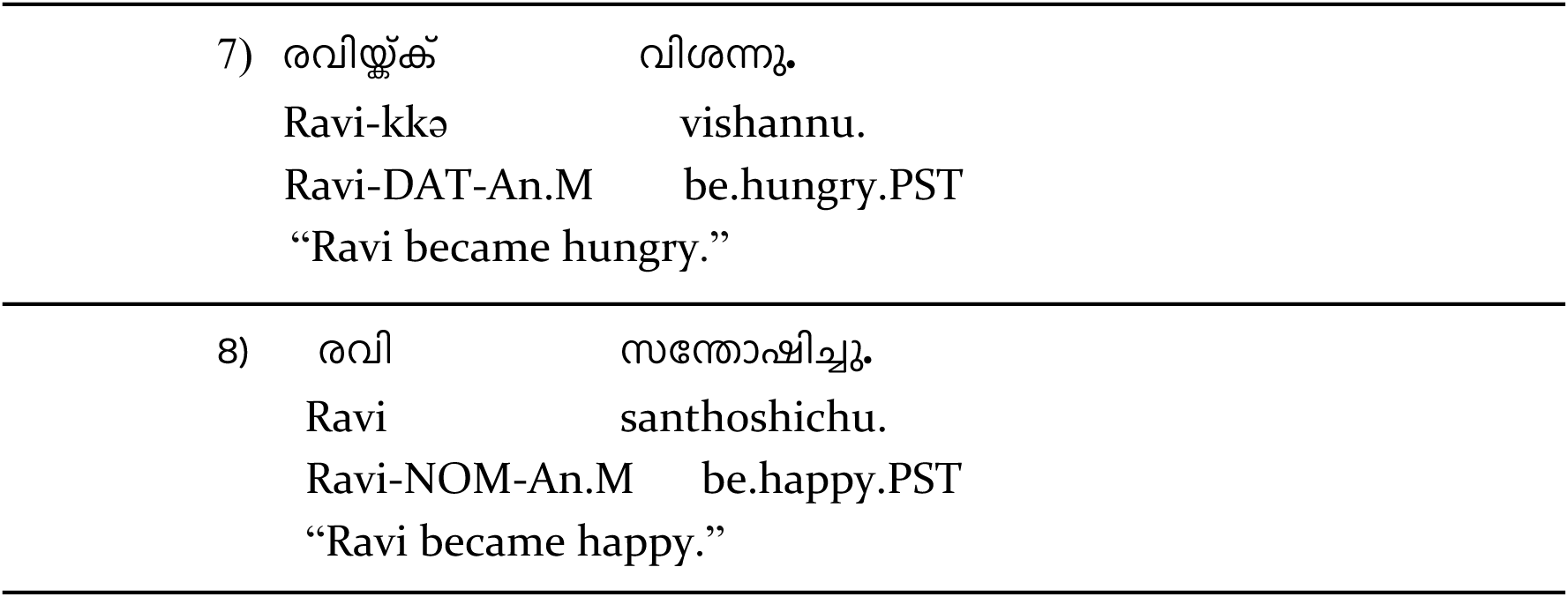

|  |  |
| --- | --- |
| 7) രവിയ്ക്ക് | വിശന്നു. |
| Ravi-kkə | vishannu. |
| Ravi-DAT-An.M | be.hungry.PST |
| “Ravi became hungry.” |  |

|  |  |  |
| --- | --- | --- |
| 8) | രവി | സന്തോഷിച്ചു. |
|  | Ravi | santhoshichu. |
|  | Ravi-NOM-An.M | be.happy.PST |
|  | “Ravi became happy.” |  |

As can be observed from examples (7) and (8), verbs pertaining to physical experiences such as “*vishakkuka* - be hungry” require a dative subject in the simple intransitive construction, whereas verbs associated with a mental experience like “*santhoshikkuka* - be happy” require a nominative subject (Mohanan, 1990; Jayaseelan, 2004; Nizar, 2010; Nair, 2012; Krishnan, 2019). The present study investigated the processing of the two types of experiencer verbs such as in (7) and (8) to study whether they are processed similarly or differently, given their varying morphosyntactic realization in Malayalam in spite of the fact that both are subject experiencer verbs.

The remainder of the paper is organized as follows. Section 2 provides a brief account of the relevance of studying experiencer verbs in Malayalam. Section 3 reports the motivation behind the present study and our hypothesis. Then, the material used in the study, participant information, and the procedure are presented in Section 4. The EEG recording, pre-processing, statistical analysis, the behavioural and ERP results are presented in Section 5. Section 6 provides a detailed discussion and interpretation of the data, and Section 7 concludes the paper.

## 2. Experiencer Verbs in Malayalam

Malayalam is a nominative-accusative South Dravidian language with agglutinative morphology, and predominantly exhibits SOV word order. However, the word order is relatively flexible, with grammatical relations and semantic roles indicated in Malayalam through a series of case suffixes and/or postpositions as in other Dravidian languages. Thus, changing the word order in a sentence typically does not change its meaning, as the roles and relations are primarily conveyed through these suffixes. Malayalam verbs do not show agreement, unlike other sister languages in the Dravidian language family, and can be categorized into various types, with one significant classification being experiencer verbs or psych verbs (Jayan and Kumar, 2018). These verbs encompass semantic notions related to experiencing, wanting, feeling, liking etc. Experiencer verbs in Malayalam are further divided into two categories based on their usage within the language: simple verbs (as in 9 and 10) and complex predicates (as in 11 and 12) (Nizar, 2010).

|  |  |  |  |
| --- | --- | --- | --- |
| 9) | അവൾക്ക് | വിശന്നു. |  |
|  | Avalkkə | vishannu. |  |
|  | She. F. Dat | be.hungry.PST. |  |
|  | “She became hungry.” |  |  |
| 10) | അവൾ | സന്തോഷിച്ചു. |  |
|  | Aval | santhoshichu. |  |
|  | She. F. Nom | be.happy.PST. |  |
|  | “She became happy.” |  |  |
| 11) | അവൾക്ക് | വിശപ്പ് വന്നു. |  |
|  | Avalkkə | vishappu vannu. |  |
|  | She. F. Dat | hunger come.PST. |  |
|  | “She became hungry.” |  |  |
| 12) | അവൾക്ക് | സന്തോഷം വന്നു. |  |
|  | Avalkkə | santhosham vannu. |  |
|  | She.F. Dat | happiness come.PST. |  |
|  | “She became happy.” |  |  |
|  |  |  | (Krishnan, 2019) |

As illustrated by the examples in (9) and (10), simple experiencer verbs in Malayalam require different cases on the subject argument based on the kind of experience the verb indicates (Jayaseelan, 2004). The complex predicate constructions in (11) and (12), on the other hand, involve predicates with two elements: the first element can be a noun, adjective, or non-finite verb, and the second element is a verb lacking semantic content known as a ‘light verb’ (Mohanan and Mohanan, 1990). When these light verbs combine with a noun, adjective, or non-finite verb, the resulting complex predicate triggers the dative case marking on the logical subject. In (11) and (12), the first element in the complex predicate, which carries the semantic meaning of experience, is morphologically related to the simple experiencer verbs in sentences (9) and (10) respectively. In sum, physical experiencer verbs require the subject argument to be in the dative case, in both their simple form and corresponding complex predicate form. By contrast, mental experiencer verbs require the subject to be in the nominative case in their simple form, and in the dative case in complex predicates (Nizar, 2010; Mohanan and Mohanan, 1990; Krishnan, 2019; Jayaseelan, 2004).

Theoretical investigations on Mental and Physical experiencer verbs in Malayalam have shown that they differ from each other in terms of both syntax and semantics. According to Mohanan and Mohanan (1990), who aimed to unify dative-induced meanings in Malayalam using a semantic role-based approach, mental experiencer verbs denote an experience of change of state in an individual (plain experiencers), while physical experiencer verbs denote an experience of advent of a new state to an individual (dative experiencers as experiencer goals). Nizar (2010), discussing the arguments of Mohanan and Mohanan (1990), explained that the individual is the goal of the advent of a state with verbs of physical experience, and thus the subject should be marked with dative case. Because the experiencer of a mental experience is not a goal, it is not marked dative, but rather is in the nominative case. Jayaseelan (2004), while discussing the subjecthood status of dative nominals, argued that mental experiencer verbs and physical experiencer verbs differ syntactically based on the presence or absence of pleonastic PRO. He also argued that the nominative subject of mental experiencer verbs indicates a greater degree of volition or control compared to the dative marked argument of physical experiencer verbs. We consider the dative marked argument of physical experiencer verbs as the subject following Nizar (2010) and Krishnan (2019). More importantly for present purposes, Krishnan (2019) builds upon Butt’s (2006) account of Urdu case alterations and applies it to Malayalam, a non-ergative language, and suggests that Malayalam employs the nominative versus dative case alteration to indicate semantic differences: more control vs less control, respectively. These differences are triggered by the predicate via the theta role and the inherent/structural case assigned to the subject. Krishnan (2019) used experiencer verbs such as “*ishttapeduka* - like” and “*dhaahikkuka* - be thirsty” cited in Jayaseelan (2004) as examples to illustrate the presence versus absence of volitionality/agency of a subject when it is marked nominative versus dative respectively. When the subject is marked dative in Malayalam, it signals the fact that the state of affairs described by the predicate is not volitional, and the subject lacks control over this state. For instance, the nominative experiencer argument in examples (13) and (16) has more control over the action, and therefore implies volition/agency, compared to the dative experiencer argument in examples (14) and (15).

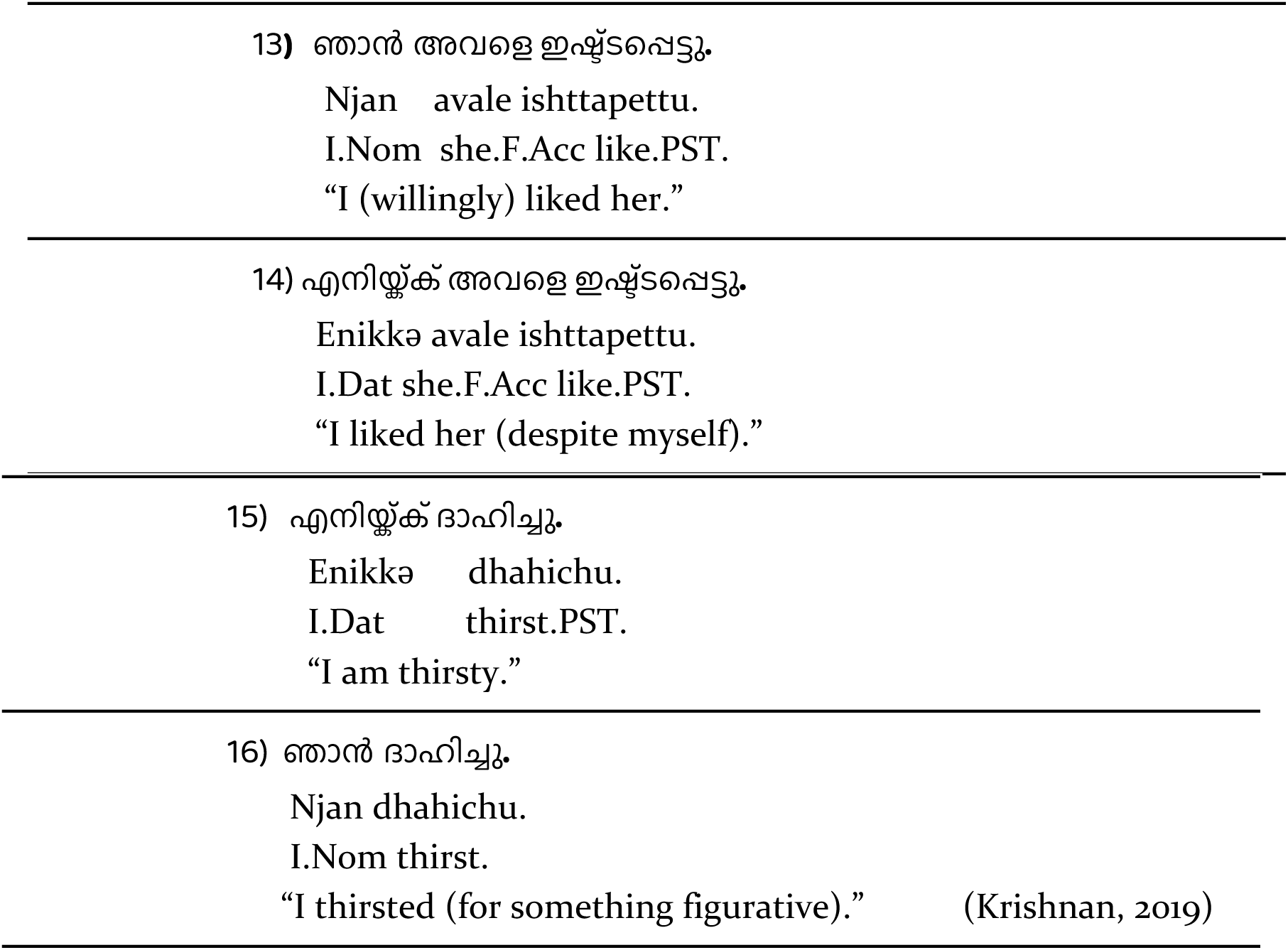

However, note that examples (14) and (16) are exceptions rather than the rule. Mental experiencer verbs pattern together syntactically with agentive verbs, and they both require nominative subjects, because the subjects of these verb types have more control/agency over the action/state. By contrast, physical experiencer verbs in their literal meaning require dative subjects. Krishnan (2019) illustrated this with an adverb test: verbs that require nominative subjects, such as mental experiencer verbs (17) and agentive verbs (19) can be modified with adverbs such as “unknowingly”, “knowingly”, whereas modifying physical experiencer verbs that require dative subjects with such adverbs results in ungrammaticality (18).

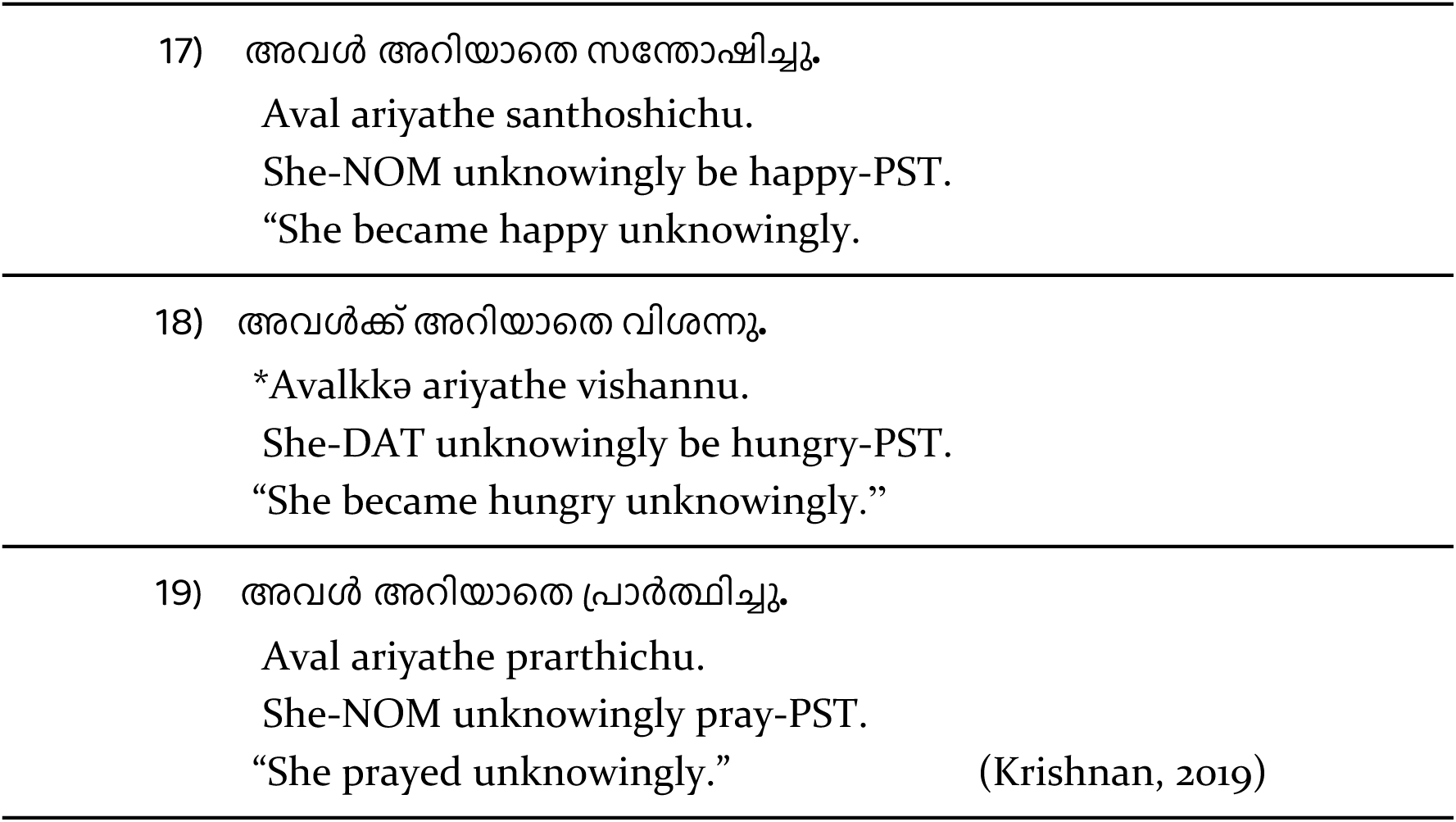

In view of this, the present study aimed to investigate the neurophysiological processing correlates of the two types of experiencer verbs in Malayalam.

## 3. The Present Study

The main goal of the current study, as mentioned in the previous sections, is to investigate whether mental and physical experiencer verbs are processed similarly, i.e. whether their neurophysiological correlates are qualitatively similar or alternatively, there are processing differences between the two.

The study employed simple intransitive subject experiencer constructions because they offer unique opportunities for Dative-Nominative alteration depending on the type of experience conveyed by mental versus physical experiencer verbs. By considering the aim of the study and structure of the experiencer verbs in Malayalam, we devised a 2*2 factorial design. We manipulated the case marking of the subject argument (i.e., Nominative/Dative) and verb type, that is, mental and physical experiencer verbs. Speakers whose first-language is Malayalam read intransitive sentences comprising dative subjects (“*Ravikkə* – Ravi-DAT”/ “*Meenuvinə* – Meenu-DAT”) or nominative subjects (“Ravi-NOM / Meenu-NOM”) followed by mental (“*santhoshichu* - be.happy”) or physical experiencer verbs (“*vishannu* - be.hungry”).

Except for a handful of studies conducted on other Dravidian languages, including Muralikrishnan (2007; 2011; 2024) and Muralikrishnan et al., (2015) on Tamil and Niharika et al., (2020) on Kannada, there is no published ERP data on Malayalam to date. We have therefore considered the theoretical perspectives on mental and physical experiencer verbs in Malayalam, as discussed in the previous section, as well as the results from existing processing literature on various types of experiencer verbs while formulating the hypotheses for our study. We hypothesize that violation conditions involving both mental and physical experiencer verbs should elicit qualitatively similar ERP correlates compared to their correct counterparts. This is in line with the theoretical literature on experiencer verbs, as both mental and physical experiencer verbs have the same argument structure (Subject_EXP_–Verb_MENT/PHY_) (Belletti and Rizzi, 1988; Grimshaw, 1990). Alternatively, if mental and physical experiencer verbs are processed differently in Malayalam, then we should observe neurophysiological differences between these two conditions. The alternative hypothesis stems from the differences in the realization of experiencer arguments. The experiencer verbs impose different morphosyntactic constraints on their experiencer arguments in Malayalam. Mental experiencer verbs require experiencer arguments to be in the nominative case, while physical experiencer verbs require dative case marking on the experiencer argument. When these case marking requirements are not met, it can lead to processing difficulties. In terms of ERP components, we expected an N400 effect for violations involving mental as well as physical experiencer verbs (Friederici and Frisch, 2000; Dröge, Maffongelli, & Bornkessel-Schlesewsky, 2014; Gattei, Tabullo, París, & Wainselboim, 2015), resulting from the conflict between the lexical-semantic information of the verb and the case information of the subject noun. In line with findings from Bornkessel et al. (2002, 2003), we further expected to observe an early positivity effect for the non-anomalous condition with mental experiencer verbs and nominative subjects, as a marker of reflecting the reanalysis from the initially preferred actor role for the nominative noun towards an experiencer role based on the verb category information.

Additionally, we hypothesize that processing physical experiencer verbs will be more costly than processing mental experiencer verbs. There are at least three motivations for this hypothesis. First, a nominative noun in the sentence-initial position entails multiple possible continuations (Intransitive, transitive and di-transitive constructions) compared to a dative case marked noun in the sentence-initial position, with which very few unmarked or felicitous continuations are possible in the language. Second, a nominative noun would minimally predict an intransitive construction with an action verb based on the minimality principle (Bornkessel and Schlesewsky, 2006; Bornkessel-Schlesewsky and Schlewesky, 2009), whereas a dative case marked noun would predict an experiencer verb, as cross-linguistic studies show that dative subjects are more commonly associated with experiencer verbs (Van Valin, 1991). That is, a nominative noun typically aligns with agents performing an action and thus would lead to an expectation for an action verb. When the parser encounters a physical experiencer verb following a nominative noun, this deviation from the expectation for an action verb would lead to increased processing costs. In contrast, a dative noun would predict for an experiencer verb, and when a mental experiencer verb (anomalous) is encountered instead of a physical experiencer verb, this would be less costly, because it still aligns with the more general expectation for an experiencer verb. Third, physical experiencer verbs indicate the advent of a new state whereas mental experiencer verbs indicate a change of state (Mohanan & Mohanan, 1990). Whilst a straightforward possibility is that there could be amplitude differences in the negativity effect for physical experiencer verbs compared to mental experiencer verbs, whether and in what manner the predicted difference between the physical and mental experiencer violation conditions is reflected in the ERPs remains to be seen.

## 4. Materials and Participants

### Ethical Statement

The research protocol for this experiment was approved by the Institute Ethical Committee (IEC) of the Indian Institute of Technology Ropar, where the experiment was conducted. Informed and written consent was obtained from participants when they arrived at the lab. All the assessments thus carried out were in accordance with the guidelines and regulations approved by the committee.

### Participants

Twenty-eight first-language speakers of Malayalam (mean age = 27.28; age range = 18–40; 11 female; 17 male), mostly students and staff at the Indian Institute of Technology Ropar, living in Ropar, India, participated in the experiment. Each participant was remunerated for their participation as per the allowance permitted by the ethical committee. Before the start of the experiment, the participants were briefed about the experiment and informed consent was obtained from them for the use of their data for academic purposes. All the participants were right-handed as evaluated by an adapted version of the Edinburgh Handedness Inventory (Oldfield, 1971) in Malayalam. The participants had normal or corrected-to-normal vision and had no known neurological disorder at the time of their participation in the experiment. All the participants were first-language speakers of Malayalam and reported having acquired the language before the age of six. As is the case with a sizeable part of the Indian population, most of our participants spoke additional languages. Data from 7 further participants were not included for analysis due to excessive artefacts.

### Materials

We employed 9 mental experiencer verbs and 9 physical experiencer verbs for constructing the critical conditions in our study. Each verb was repeated 4 times with 4 different proper nouns (2 masculine and 2 feminine nouns) which resulted in 36 sets of sentences in four critical conditions as in Table 1, thus yielding 144 critical sentences. The noun at NP was either nominative (null case marking) or dative marked. Since Malayalam dative case marker has two allomorphs “-kkə” and “-ə”^1^, we used an equal number of nouns requiring either the “-kkə” or the “-ə” marker (18 each). Of the 144 critical sentences, 72 sentences were grammatically correct (half of them with dative subjects and the other half with nominative subjects) and 72 sentences were grammatically incorrect (half of them with dative subjects and the other half with nominative subjects). Additionally, we constructed 288 filler items to introduce variety in the stimuli to avoid strategic responses from the participants. These fillers were interspersed with the critical items such that there was a total of 432 sentences. These items were pseudorandomized for presentation during the experiment.

**Table 1.** The example sentences for the present study. (Experiencer verbs are expressed as MENT (mental experiencer) and PHY (physical experiencer) and case markers are expressed as NOM (nominative) and DAT (Dative), NOM-MENT and DAT-PHY are grammatically correct sentences, DAT-MENT and NOM-PHY are ungrammatical ones).

| Condition | Sample Stimulus |
| --- | --- |
| NOM-MENT | രവി സന്തോഷിച്ചു.<br>Ravi santhoshichu.<br>Ravi-NOM-An.M be.happy.PST<br>“Ravi became happy.” |
| DAT-MENT | *രവിക്ക് സന്തോഷിച്ചു.<br>Ravi-kkə santhoshichu.<br>Ravi-DAT-An.M be.happy.PST<br>“Ravi became happy.” |
| NOM-PHY | * രവി വിശന്നു.<br>Ravi vishannu.<br>Ravi-NOM-An.M be.hungry.PST<br>“Ravi became hungry.” |
| DAT-PHY | രവിക്ക് വിശന്നു.<br>Ravi-kkə vishannu.<br>Ravi-DAT-An.M be.hungry.PST<br>“Ravi became hungry.” |

### Procedure

The experiment consisted of a practice session followed by the actual experiment, with all the activities, including electrode preparation and stimulus presentation lasting approximately 3 hours. Initially, the procedure and the tasks to be performed during the experiment were explained to the participants, and a printed instruction sheet was provided to them. However, the question under investigation in the experiment was not revealed to the participants for the acquisition of unbiased data. Then, they filled a consent form giving informed consent for their participation along with a Malayalam version of the Edinburg handedness questionnaire to evaluate their handedness. Finally, the head measurements of the participants were taken, and the Hydrocel GSN net was placed on their scalp. They were seated within a soundproof chamber on a comfortable chair at a distance of 1 m from a 20” LCD monitor on which the stimuli were presented using E-prime 2.0 (Psychology Software Tools, Pittsburgh, PA) (https://pstnet.com).

The structure of each experimental trial was as follows. Each trial commenced with a fixation ‘+’ sign at the centre of the screen for a period of 1000 ms. After a blank screen for 100 ms, the rapid serial visual presentation of the stimulus sentence commenced, consisting of two separate chunks for critical as well as filler trials. Each of the chunks was presented for a period of 650 ms followed by an inter-stimulus interval (ISI) for a duration of 100 ms. This presentation rate provided a comfortable reading pace for participants and is in line with several existing studies on languages that are orthographically and/or morphologically complex, such as Arabic (Muralikrishnan & Idrissi, 2021), Mandarin Chinese (Wang et al., 2009), Icelandic (Bornkessel-Schlesewsky et al., 2020), Japanese (Wang & Schumacher, 2013) and Turkish (Demiral et al., 2008).

After the last chunk, participants performed two tasks. As a violation paradigm was used in the study, they performed an acceptability judgment task after the stimulus was displayed whereby, they had to press a green button on a response pad if they found the sentence displayed earlier to be acceptable or a red button if they found the sentence to be unacceptable. As a cue to the acceptability judgment task, a “???” sign appeared on the screen after the last chunk. In addition, to ensure that they read the sentence attentively, the participants performed a probe task. After finishing the acceptability judgment task or when 1500 ms had passed, a probe word appeared on the screen for 2000 ms. Participants were then required to press the green button if the probe word was present in the preceding sentence, or the red button if it was not. Half of the probe words were present in the preceding trial and required a ‘yes’ response, whereas the other half were novel and required a ‘no’ response.

The placement of the green and red buttons was counterbalanced across participants such that, the ‘yes’ response was associated with the left key and the ‘no’ response was associated with the right key for half the participants, and vice versa for the other half. Participants were requested to avoid blinks during the presentation of the words, but they could blink while performing the two tasks.

Before the actual experiment, they were given a practice session with 12 items to familiarize themselves with the experimental trial structure. None of the actual experimental stimuli occurred in the practice session. The experimental session was subdivided into 9 blocks of 48 sentences each, with short breaks at the end of each block. At the end of the experiment, the participants were requested to fill a questionnaire regarding their experience with the experiment.

## 5. EEG recording, pre-processing and statistical analysis

The EEG was recorded by means of 32 Ag/AgCI electrodes fixed at the scalp using the Hydrocel Geodesic Sensor Net, with Cz serving as the online reference. The electrooculogram was recorded for monitoring horizontal and vertical eye movements using electrodes placed at the outer canthi and under the eyes respectively, and the interelectrode impedance was kept below 50 kΩ (amplifier input impedance > 1 GOhm) as per system recommendations (Ferree et al., 2001). All EEG and EOG signals were amplified using a Net Amps 400 Amplifier. The data was recorded at a sampling rate of 500 Hz.

The EEG data thus recorded was pre-processed for further analysis using the EEGLAB toolbox (Version 14; Delorme & Makeig, 2004, sccn.ucsd.edu) in MATLAB (Version R2023b; The MathWorks, Inc.). The data was down-sampled to 250 Hz and filtered using a 0.3–20 Hz band-pass filter in order to remove slow signal drifts. These filter settings are sufficiently broad to include language-related ERP activity that is typically in the frequency range of about 0.5–5 Hz (Delorme, 2023; Roehm, Winkler, Swaab & Klimesch, 2002), and have been employed in several previous cross-linguistic ERP studies on language processing in diverse languages. This data was then re-referenced offline to the average of the two mastoids, and channels on the outer extent of the face and head were removed. In a 1 Hz high-pass filtered copy of this original data, an Independent Component Analysis (ICA, Iriarte et al., 2003) was computed. Bad channels were removed in this data before submitting it to an ICA computation using the extended Infomax algorithm. The resulting Independent Components (ICs) were then tested using the ICLabel plugin (Pion-Tonachini, Kreutz-Delgado & Makeig, 2019) for identifying and marking ICs that were artefactual. The weights computed during ICA were then copied to the original data, and the ICs marked as artefactual were rejected from the original data. The bad channels removed before computing the ICA were interpolated from the remaining channels. The data was then imported in R (Version 4.4.2; R Core Team, 2024) using the eeguana package (Version 0.1.11.9001; Nicenboim, 2018) for epoching and statistical analysis.

Data epochs were extracted from the continuous data for each participant for the critical conditions in the critical position, i.e., the verb, from 200 ms before the onset until 1200 ms after the onset (i.e., -200 to 1200 ms). Epochs in which the amplitude exceeded the threshold of 100 μV in either direction were rejected, as well as those in which the difference between the minimum and maximum amplitudes within a window of 200 ms crossed the threshold of 100 μV. Furthermore, trials in which the acceptability judgement task was not performed were also rejected. Data from participants with too few remaining trials were excluded from further analysis. For a given participant, data epochs from about 27 to 28 trials remained in each condition after these rejections, and therefore the number of valid trials entering the analysis did not differ much across conditions. Across participants, there were a total of 3228 data epochs / trials entering the analysis, with about 800 trials per critical condition. For the purposes of visualization, the valid epochs were averaged across items per condition per participant, and then the grand-averages were computed across participants and smoothed using an 8 Hz low-pass filter to obtain ERP plots at the verb for each condition.

ERP Data Analysis. The single trial EEG epochs at the NP and the verb for each critical condition were used for statistically analysing the mean amplitudes in selected time-windows of interest by fitting linear mixed effects models using the lme4 package (Bates et al., 2015) in R (Version 4.4.2, R Core Team 2024). The statistical models included the fixed factors Case (Nominative vs Dative) and Verb type (Mental experiencer vs Physical experiencer), as well as the topographical factor Regions of Interest (ROI). The ROIs were defined by clustering topographically adjacent electrodes in 6 lateral and 2 midline regions. The lateral ROIs were: Left-Frontal, comprised of the electrodes E3 and E11 (equivalent to F3 and F7 in the 10-20 electrode system); Left-Central, comprised of the electrodes E5 and E13 (C3 and T7); Left-Parietal, comprised of the electrodes E7 and E15 (P3 and P7); Right-Frontal, comprised of the electrodes E4 and E12 (F4 and F8); Right-Central, comprised of the electrodes E6 and E14 (C4 and T8); and Right-Parietal, comprised of the electrodes E8 and E16 (P4 and P8). The midline ROIs were: Mid-Fronto-Central, comprised of E17 and E28 (Fz and ∼FCz); and Mid-Parieto-Occipital, comprised of E19, E20, E9 and E10 (Pz, Oz, O1 and O2). Furthermore, rather than performing a traditional subtraction-based baseline correction, the mean 200 ms pre-stimulus baseline amplitude (-200 to 0 ms) in each data epoch was included as a covariate (after scaling) in the statistical model. This was done in order to account for and regress out potential contributions of differences in the baseline period in the statistical analysis (Alday, 2019). However, we do not interpret effects involving prestimulus amplitudes, because these did not form part of our hypotheses. Further, this is in line with the fact that “we can include additional covariates as controls without further interpreting those covariates” (Alday, 2019, p.9). The contrasts for the categorical factors used scaled sum contrasts (effects coding), such that coefficients reflect differences to the grand mean (Schad et al., 2020). Except where specified otherwise, the random effects structure was maximal (Barr et al., 2013), with random intercepts for participants and items, and by-participant random slopes for the effect of Verb type, Case and their interaction term. A by-item random slope specification was not included because it would not be identifiable for this data: the items were coded uniquely across conditions such that they indicated a combination of condition + item. Following modern statistical recommendations, we do not use the term ‘statistically significant’ or its variants based on p-value thresholds (Wasserstein et al., 2019), but report precise p-values as continuous quantities (e.g., p = 0.06 rather than p < 0.08), unless a value is “below the limit of numerical accuracy of the data”, in which case, we report it as p < 0.001 (Amrhein et al., 2019, p.206). Further, we supplement the p-values by transforming them into s-values (Shannon information, surprisal, or binary logworth) and report s = – log_2_(p), which provides a nonprobability measure of the information provided by a p-value on an absolute scale (Shannon, 1948; Greenland, 2019). In other words, “the s-value provides a gauge of the information supplied by a statistical test” and has the advantage of providing “a direct quantification of information without” requiring prior distributions as input (Rafi & Greenland, 2020, p.6).

## Results

### Behavioural data

The mean acceptability ratings as well as the probe detection accuracy for the critical conditions, shown in Table 2, were calculated for the trials that entered the ERP analysis using the behavioural data collected during the experiment. As mentioned earlier, only those trials in which the acceptability judgement task was performed (i.e., not timed out) were considered for analysis. Acceptability was highest for the grammatical conditions, whereas it was lowest for the violation conditions. The probe detection accuracy was very high across all conditions. Fig. 1 shows raincloud plots (Allen, Poggiali, Whitaker, Marshall, & Kievit, 2021) of the behavioural acceptability judgements. Panel A shows the by-participant variability of acceptability ratings, with the individual data points representing the mean by-participant acceptability of each case and verb type combination. Panel B shows the by-item variability of acceptability ratings, with the individual data points representing the mean by-item acceptability of each case and verb type combination.

**Fig. 1.**
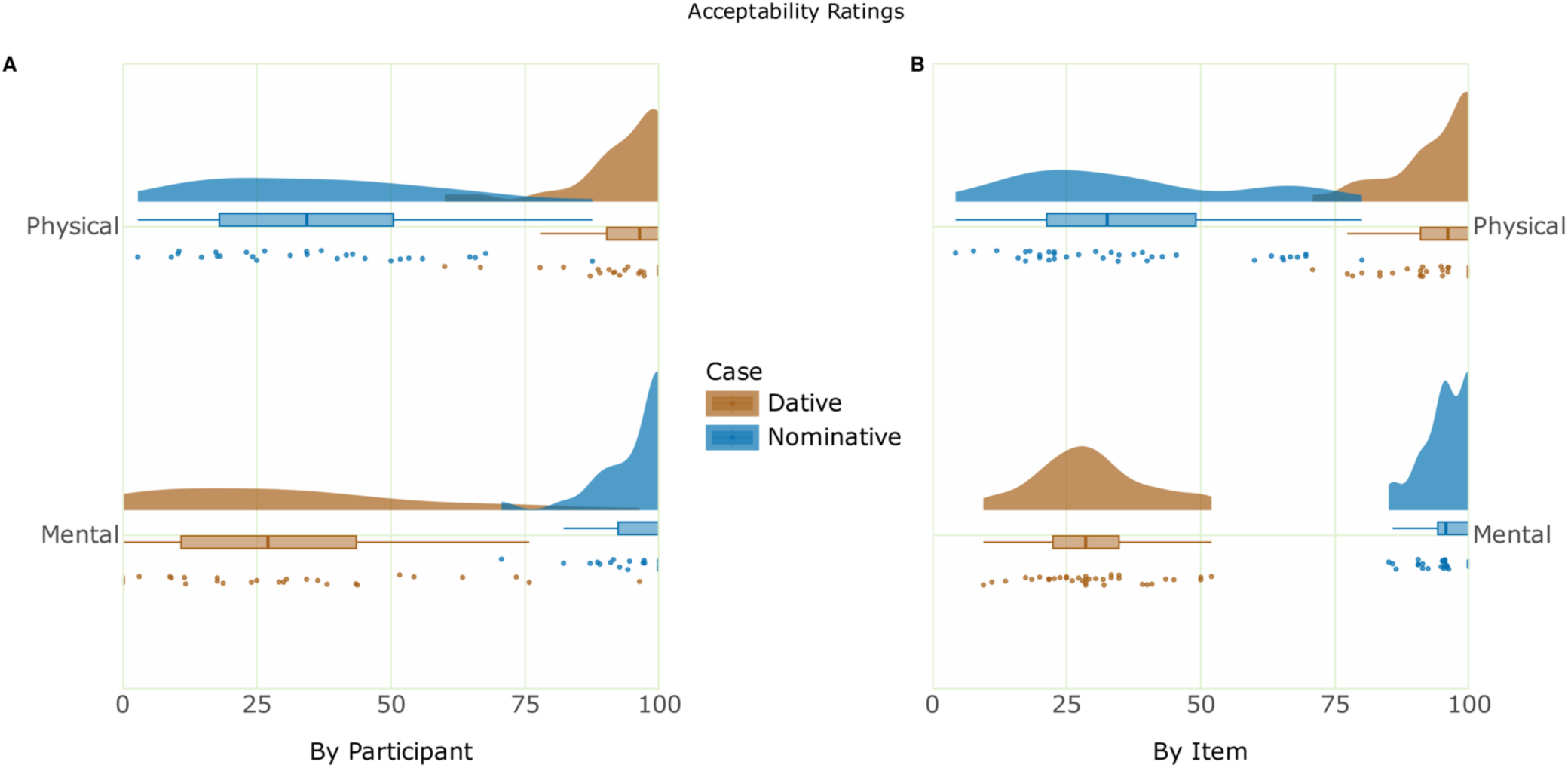
Raincloud plot of the acceptability ratings, showing by-participant variability (Panel A) and by-item variability (Panel B).

**Table 2.** (Mean acceptability ratings and probe detection accuracy)

| Case | Verb type | Mean Acceptability % (SD) | Mean Accuracy % (SD) |
| --- | --- | --- | --- |
| Nom | Mental | 95.7 (20.3) | 99.0 (10.0) |
| Dat | Mental | 29.8 (45.8) | 95.5 (20.8) |
| Nom | Physical | 36.4 (48.1) | 95.4 (20.9) |
| Dat | Physical | 93.4 (24.9) | 98.5 (12.2) |

The behavioural acceptability and accuracy were analysed by fitting generalised linear mixed models using the lme4 package (Bates et al., 2015) in R. In the analysis of acceptability data, the statistical models included the fixed factors Case (Nominative vs Dative) and Verb type (Mental experiencer vs Physical experiencer), with random intercepts for participants and items, and by-participant random slopes for the effect of Case, Verb type and their interaction term. Type II Wald chi-square tests of the fitted model (AIC = 2352.56) of the acceptability data showed an interaction of Verb x Case (χ^2^(1) = 111.57, p < 0.001, s = 84.22) that was more informative than main effects of the individual factors Verb (χ^2^(1) = 0.08, p = 0.77, s = 0.36) and Case (χ^2^(1) = 5.94, p = 0.01, s = 6.08). Estimated marginal means on the response scale were computed on the model using the emmeans package (Lenth, 2021) to resolve this interaction, which showed that the estimate for mental experiencer verbs with dative subjects was less (estimate = 0.22, SE = 0.069, p = 0.001, s = 9.65) than that for mental experiencer verbs with nominative subjects (estimate = 0.97, SE = 0.009, p = 0, s = ∞). By contrast, the estimate for physical experiencer verbs with nominative subjects was less (estimate = 0.31, SE = 0.059, p < 0.001, s = 23.28) compared to that for physical experiencer verbs with dative subjects (estimate = 0.95, SE = 0.013, p = 0, s = ∞). The pairwise contrasts of these estimates within each level of Verb type showed that, the contrast between the subject types for mental experiencer verbs (estimate = 0.749, SE = 0.072, p < 0.001, s = 81.75) was similarly highly informative to that for physical experiencer verbs (estimate = -0.640, SE = 0.060, p < 0.001, s = 84.64). This is a reflection of the very low acceptability for violation conditions versus their non-anomalous counterparts regardless of the verb type.

In the analysis of the probe detection accuracy, models with an interaction term of the fixed factors Verb and Case, as well as those with random slopes were singular. The model with Verb and Case as fixed factors with by-participant and by-item random intercepts (AIC = 846.29) showed no effects of Verb or Case (s < 1), reflecting the highly similar (ceiling) accuracy of probe detection across conditions.

### ERP data

The ERPs at the critical verb position are shown in **Fig. 2**. The ERPs at the sentence-initial noun are not part of our hypotheses, but are provided Fig. S1 in the supplementary material. Visual inspection of the ERP data showed that the violation conditions engendered a negativity effect at the verb compared to their non-anomalous counterparts. In addition to considering ERP components that are relevant for language processing based on previous literature, this qualitative observation based on visual inspection informed the choice of time-window for analysis, namely 400–800 ms. This is in line with a similarly long time-window selected for analysis in a previous study on Turkish, with the critical position for analysis also at the position of the sentence-final verb (Bornkessel et al., 2011). Besides this similarity to our study, Turkish is also a verb-final language which is morphologically rich and similar (agglutinative) to Malayalam. The single trial ERP mean amplitudes extracted in the analysis time-window from a total of 3228 data epochs / trials entered the analysis as mentioned earlier, with about a median of 800 trials per critical condition. The raw data collected during the experiment, the pre-processing pipeline used and the pre-processed data are available in the data respository online.

**Fig. 2.**
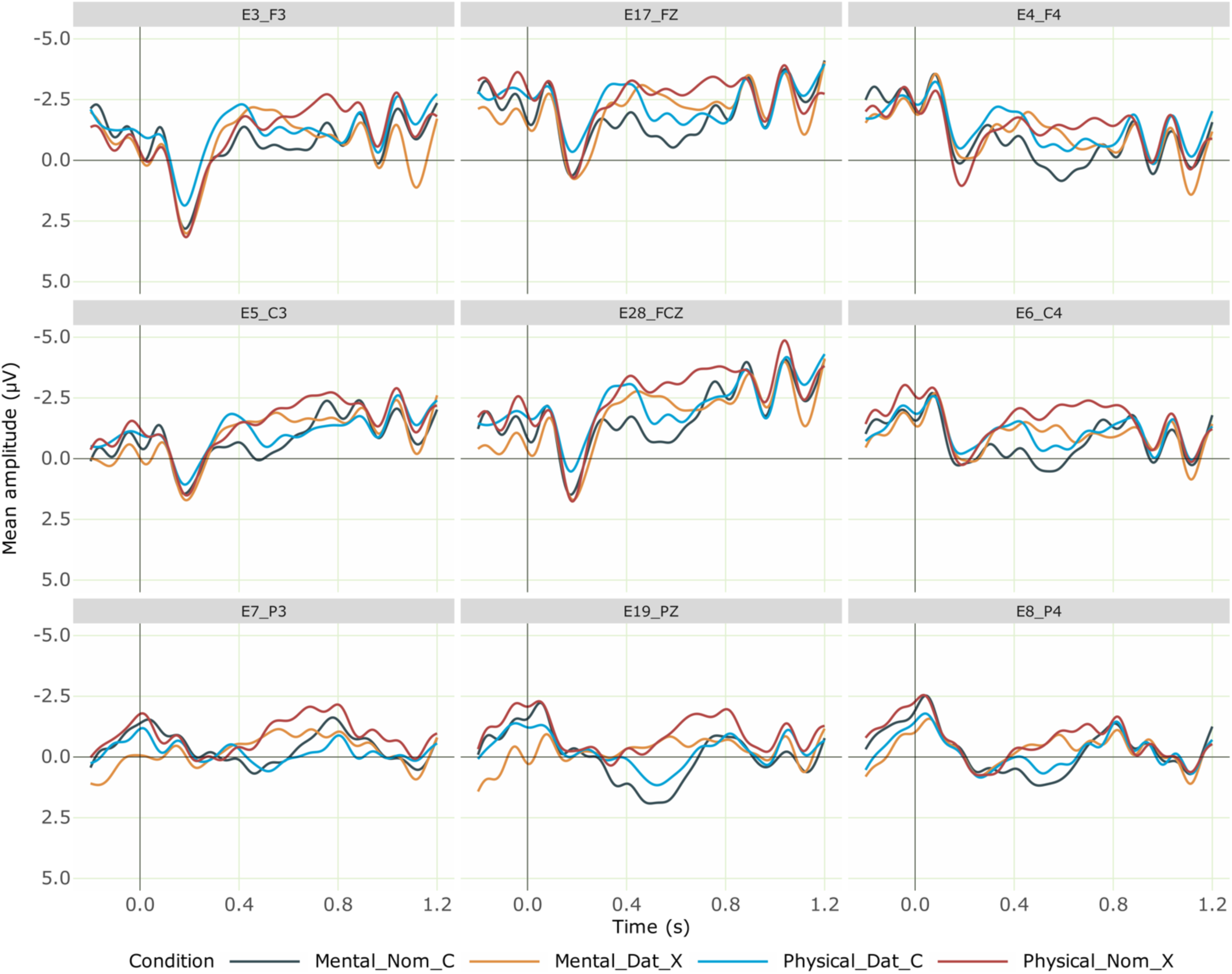
Grand averaged ERPs at the verb for the critical conditions from 28 participants. Negativity is plotted upwards; the time axis runs from -0.2 s to 1.2 s (i.e., -200 ms to 1200 ms) with 0 being the onset of the critical verb. The dark blue line represents the correct mental experiencer verb (with a nominative subject), while the orange line shows the incorrect mental experiencer verb (with a dative subject), which elicited a negativity effect. The light blue line indicates the correct physical experiencer verb (with a dative subject), and the red line represents the incorrect physical experiencer verb (with a nominative subject), which elicited a negativity effect.

### Planned Analysis

We first computed a linear mixed effects model m0 involving the fixed factors Verb type, Case of the subject noun, ROI, the covariate PrestimMeanAmp (centred), and by-participant and by-item random intercepts. The analysis code and full model outputs are available as R notebooks in the analysis repository online. Type II Wald Chi-squared tests on this model (AIC = 428205.60) showed an interaction of Verb x Case (χ^2^(1) = 10.6, p = 0.001, s = 9.81) that was more informative than main effects of the individual factors Verb and Case (s < 1), as well as their two-way and three-way interactions with ROI (s < 1). Estimated marginal means on the response scale were computed on the model using the emmeans package (Lenth, 2021) to resolve this interaction, which showed that the estimate for mental experiencer verbs with dative subjects was more negative (estimate = -1.043, SE = 0.445, p = 0.01, s = 5.70) than that for mental experiencer verbs with nominative subjects (estimate = -0.093, SE = 0.445, p = 0.83, s = 0.26). By contrast, the estimate for physical experiencer verbs with nominative subjects was more negative (estimate = -1.076, SE = 0.445, p = 0.01, s = 6) compared to that for physical experiencer verbs with dative subjects (estimate = -0.346, SE = 0.445, p = 0.43, s = 1.20). The pairwise contrasts of these estimates within each level of Verb type showed that, the contrast between the subject types for mental experiencer verbs (estimate = 0.949, SE = 0.372, p = 0.01, s = 6.52) was similar to, but slightly more informative than, that for physical experiencer verbs (estimate = -0.729, SE = 0.372, p = 0.04, s = 4.32). This pattern suggests that there is statistical support to interpret the ERPs for the violation conditions as more negative (broadly) across the scalp than their respective non-anomalous counterparts.

In order to verify if the model fit could be improved, we then computed another model m1 by making the random effects structure in m0 maximal, namely by including by-participant random slopes for the factors Verb, Case and their interaction term. As mentioned earlier, a by-item random intercept specification is not identifiable for our data in view of how items were coded in the data, and therefore the by-item random intercept specification in m0 was retained as is in m1. Such a model would better account for within-participant ERP variance between conditions. Indeed, the model fit showed an improvement (AIC = 427161.14), and type II Wald Chi-squared tests on this improved model again showed that the interaction of Verb x Case remained more informative (χ^2^(1) = 3.56, p = 0.05, s = 4.07) than main effects of the individual factors Verb and Case (s < 1), as well as their two-way and three-way interactions with ROI (s < 1). Estimated marginal means on the response scale computed on model m1 to resolve this interaction showed that, as in model m0, the estimate for mental experiencer verbs with dative subjects was more negative (estimate = -1.243, SE = 0.569, p = 0.02, s = 5.11) than that for mental experiencer verbs with nominative subjects (estimate = - 0.024, SE = 0.595, p = 0.96, s = 0.04). Again, as in model m0, the estimate for physical experiencer verbs with nominative subjects was more negative (estimate = -1.126, SE = 0.490, p = 0.02, s = 5.53) compared to that for physical experiencer verbs with dative subjects (estimate = -0.254, SE = 0.541, p = 0.63, s = 0.64). The pairwise contrasts of these estimates within each level of Verb type showed that, the contrast between the subject types for mental experiencer verbs (estimate = 1.219, SE = 0.778, p = 0.11, s = 3.09) was similar to, but in contrast to model m0, slightly less informative than, that for physical experiencers (estimate = -0.871, SE = 0.54, p = 0.10, s = 3.22). These numerical differences between the two models notwithstanding, the overall pattern of results remain intact, namely that there is a general effect of violation such that both violation conditions elicit larger negativity effects in this time-window than their respective non-anomalous counterparts.

There are however two unplanned observations that we made about the data that are worth noting here. One, the negativity effects for the two anomalous conditions visually appear to have slightly different morphologies. The negativity for the mental experiencer violation condition appears to show an early maximum, whereas the opposite is true for the physical experiencer violation condition. Two, although the behavioural acceptability ratings were similarly low for both violation conditions, as the raincloud plots of these ratings in Fig. 1 show, a stark difference is apparent between the by-participant and by-item distribution of acceptability ratings. Whereas the variance in by-participant ratings is similar for violations involving either verb type, the by-item distribution of the ratings is spread out for physical experiencer violations but concentrated on the lower end of the acceptability spectrum for the mental experiencer violations. In order to find out if there is some statistical support in the data for these observations, we conducted two post-hoc analyses, one each addressing each of these issues, which we report in the following section.

### Post-hoc Analyses

The post-hoc analyses of the data we conducted are as follows. First, in view of the slightly different morphology of the negativity effects for the two anomalous conditions, we split the original time-window for analysis reported above into two equal time-windows, namely 400–600 ms and 600–800 ms and computed linear mixed effects models similar to m1 (i.e., with by-participant random slopes and by-item random intercepts) in each of these time-windows. Second, in order to investigate if the ERPs varied contingent upon the post-trial acceptability ratings, we conducted an acceptability-contingent analysis of the single trial ERP amplitudes in the original time-window (400–800 ms) for which we reported the planned comparisons above. The analysis code and full model outputs of these analyses are available as R notebooks in the analysis repository online.

### Split-window analyses

Time-window 1: 400–600 ms. A linear mixed effects model similar to m1 with maximal random effects specification for the single-trial ERP amplitudes in this time-window revealed the following. Type II Wald Chi-squared tests on this model (AIC = 430093.97) showed an interaction of Verb x Case (χ^2^(1) = 5.031, p = 0.02, s = 5.32) that was more informative than main effects of the individual factors Verb and Case (s <= 1), as well as their two-way and three-way interactions with ROI (s <= 1). Estimated marginal means on the response scale were computed on the model to resolve this interaction, which showed that the estimate for mental experiencer verbs with dative subjects was more negative (estimate = -1.375, SE = 0.560, p = 0.01, s = 6.14) than that for mental experiencer verbs with nominative subjects (estimate = 0.302, SE = 0.539, p = 0.57, s = 0.79). By contrast, the estimate for physical experiencer verbs with nominative subjects was more negative (estimate = -0.901, SE = 0.487, p = 0.06, s = 3.95) compared to that for physical experiencer verbs with dative subjects (estimate = -0.192, SE = 0.551, p = 0.72, s = 0.46). The pairwise contrasts of these estimates within each level of Verb type showed that, the contrast between the subject types for mental experiencer verbs (estimate = 1.678, SE = 0.731, p = 0.02, s = 5.52) was more informative than that for physical experiencer verbs (estimate = -0.708, SE = 0.567, p = 0.21, s = 2.24). This pattern suggests that the negativity effect evoked by violations is of a larger magnitude in this time-window for the mental experiencer verbs compared to that for the physical experiencer verbs.

Time-window 2: 600–800 ms. A linear mixed effects model similar to m1 with maximal random effects specification for the single-trial ERP amplitudes in this time-window revealed the following. Type II Wald Chi-squared tests on this model (AIC = 437812.54) showed an interaction of Verb x Case (χ^2^(1) = 2.173, p = 0.14, s = 2.83) that was hardly more informative than main effects of the individual factors Verb and Case (s <= 1.5), as well as their two-way and three-way interactions with ROI (s <= 1). Nevertheless, estimated marginal means on the response scale were computed on the model to resolve this interaction, which showed that the estimate for mental experiencer verbs with dative subjects was more negative (estimate = - 1.110, SE = 0.610, p = 0.06, s = 3.85) than that for mental experiencer verbs with nominative subjects (estimate = -0.343, SE = 0.702, p = 0.62, s = 0.67). By contrast, the estimate for physical experiencer verbs with nominative subjects was more negative (estimate = -1.352, SE = 0.537, p = 0.01, s = 6.40) compared to that for physical experiencer verbs with dative subjects (estimate = -0.310, SE = 0.579, p = 0.59, s = 0.75). The pairwise contrasts of these estimates within each level of Verb type showed that, the contrast between the subject types for mental experiencer verbs (estimate = 0.767, SE = 0.869, p = 0.37, s = 1.40) was less informative than that for physical experiencer verbs (estimate = -1.041, SE = 0.591, p = 0.07, s = 3.67). This pattern is the opposite of what was observed in the 400–600 ms time-window and suggests that the negativity effect evoked by violations is of a larger magnitude in this time-window for the physical experiencer verbs compared to that for the mental experiencer verbs.

Taken together, the split-window analyses suggest that there is statistical support to interpret the negativity effects for the violation conditions as relatively more negative for mental experiencer verbs *earlier* compared to physical experiencer verbs, and relatively more negative for physical experiencer verbs *later* compared to mental experiencer verbs. That is, whereas the negativity effect starting at about 400 ms after the onset of the verb continues to be more negative as opposed to their non-anomalous counterparts until about 800 ms for both violation conditions, there is a difference in when the negativity peaks for the two verb types. The negativity effect reaches an early maximum by 600 ms for mental experiencer verbs, whereas it reaches a later maximum after 600 ms for physical experiencer verbs.

### Acceptability-contingent analysis

In order to examine if and in what manner the post-trial sentence acceptability possibly modulated the ERPs, we conducted an acceptability-contingent analysis. To this end, we included the sentence-final acceptability rating, coded as a factor with two levels (Acceptable and NotAcceptable; scaled sum contrasts), as a fixed effect in the linear mixed effects model m1. As mentioned earlier, the motivation for this analysis was based on the observation that there were differences apparent between the by-participant and by-item *distribution* of acceptability ratings for the two verb types. Consequently, we figured that the trial sequence might be crucial information to take into account when conducting this analysis. Therefore, the data epoch / trial number (centred) was included as a covariate so that possible changes in effects over the course of the experiment could be detected. As in model m1, all fixed factors and covariates and their interaction terms constituted the fixed effects in this new model m1a, and by-participant and by-item intercepts as well as by-participant random slopes for the factors Verb, Case and their interaction term constituted the random effects specification. In line with our motivations, we will specifically interpret interactions of Verb type and/or Case with Acceptability and Trial in this analysis.

Model m1a (AIC = 426765.99) proved to be a better description of the ERP data in the 400–800 ms time-window in comparison to the models reported in the planned analysis m0 (AIC = 428205.60) and m1 (AIC = 427161.14). Type II Wald Chi-squared tests on model m1a showed that, the main effects of Acceptability (χ^2^(1) = 4.549, p = 0.03, s = 4.92) and Trial (χ^2^(1) = 17.265, p < 0.001, s = 14.90) and the interaction Trial x Acceptability (χ^2^(1) = 33.374, p < 0.001, s = 4.92) were all informative. The interactions Verb x Trial (χ^2^(1) = 13.01, p < 0.001, s = 11.65) and Verb x Acceptability (χ^2^(1) = 8.489, p = 0.003, s = 8.129) were both very informative; the interaction Case x Trial (χ^2^(1) = 60.826, p < 0.001, s = 47.18) was much more informative than the interaction Case x Acceptability (χ^2^(1) = 2.038, p = 0.153, s = 2.70). More pertinent for present purposes, the three-way interaction Verb x Case x Trial (χ^2^(1) = 4.494, p = 0.03, s = 4.87) was relatively less informative than Verb x Case x Acceptability (χ^2^(1) = 112.01, p < 0.001 , s = 84.540), which was very highly informative; the three-way interactions Verb x Trial x Acceptability (χ^2^(1) = 60.617, p < 0.001, s = 47.03) and Case x Trial x Acceptability (χ^2^(1) = 21.946, p < 0.001 , s = 18.44) were both very informative. By contrast, the four-way interaction Verb x Case x Trial x Acceptability was not informative (χ^2^(1) = 0.306, p = 0.57, s = 0.78).

Estimated marginal means on the response scale were computed on the model to resolve the highly informative three-way interaction Verb x Case x Acceptability, which showed the following.

For mental experiencer verbs with nominative subjects (i.e., non-anomalous sentences), the estimate for sentences rated as NotAcceptable was much more negative (estimate = -4.425, SE = 0.757, p < 0.001, s = 27.55) than that for sentences rated as Acceptable (estimate = 0.153, SE = 0.602, p = 0.79, s = 0.32). By contrast, for mental experiencer verbs with dative subjects (i.e., violation sentences), the estimate for sentences rated as NotAcceptable was hardly any different (estimate = -1.288, SE = 0.552, p = 0.01, s = 5.66) from that for sentences rated as Acceptable (estimate = -1.14, SE = 0.569, p = 0.04, s = 4.47).

For physical experiencer verbs with nominative subjects (i.e., violation sentences), the estimate for sentences rated as NotAcceptable was less negative (estimate = - 0.744, SE = 0.482, p = 0.12, s = 3.02) than that for sentences rated as Acceptable (estimate = -1.580, SE = 0.494, p = 0.001, s = 9.48). By contrast, for physical experiencer verbs with dative subjects (i.e., non-anomalous sentences), the estimate for sentences rated as NotAcceptable was more negative (estimate = - 2.472, SE = 0.639, p = 0.0001, s = 13.15) than that for sentences rated as Acceptable (estimate = -0.125, SE = 0.541, p = 0.81, s = 0.29).

The pairwise contrasts of these estimates within each level of Verb type and Case showed that, the contrast between NotAcceptable and Acceptable ratings for mental experiencer verbs with (non-anomalous) nominative subjects (estimate = - 4.579, SE = 0.476, p < 0.001, s = 70.11) was much more informative than that for mental experiencer verbs with (anomalous) dative subjects (estimate = -0.146, SE = 0.214, p = 0.49, s = 1.01). By contrast, the contrast between NotAcceptable and Acceptable for physical experiencer verbs with (anomalous) nominative subjects (estimate = 0.836, SE = 0.203, p < 0.001, s = 14.57) was relatively less informative than that for physical experiencer verbs with (non-anomalous) dative subjects (estimate = -2.346, SE = 0.365, p < 0.001, s = 32.79).

Given that the estimates are on the response scale, they are indeed estimates of the mean ERP amplitude in the 400–800 ms time-window. Therefore, the pattern above suggests that, on the one hand, there is a stark difference in the ERP amplitudes for mental experiencer verbs with nominative subjects (i.e., non-anomalous sentences) when such a sentence was rated as NotAcceptable (more negative-going ERP) versus Acceptable. There is a similar, albeit numerically smaller, difference in the ERP amplitudes for physical experiencer verbs with dative subjects (i.e., non-anomalous sentences) when such a sentence was rated as NotAcceptable (more negative-going ERP) versus Acceptable. On the other, there is negligible difference in the ERP amplitudes for mental experiencer verbs with dative subjects (i.e, anomalous sentences) when such a sentence was rated as NotAcceptable versus Acceptable. Whilst there is little difference in the ERP amplitudes for physical experiencer verbs with nominative subjects (i.e., anomalous sentences) when such a sentence was rated as NotAcceptable versus Acceptable, the ERPs for Acceptable ratings are more negative, albeit numerically small, than that for NotAcceptable ratings.

In view of the additional interaction of Verb type, Case and Trial, this intriguing pattern is best examined further by resolving the higher order interaction involving all relevant factors, namely Verb type, Case, Trial and Acceptability. Estimated marginal means on the response scale computed on the acceptability-contingent analysis model m1a to resolve this interaction are plotted in Fig. 3, which illustrates the pattern described above. For each condition, the estimates (ERP amplitudes) are plotted for sentences in that condition rated as Acceptable versus NotAcceptable. Given that the ERPs preceded the acceptability ratings in time, and were engendered during the processing, the pattern of results described above in conjunction with Fig. 3 can be summarised as follows. For anomalous sentences of either verb type, there was hardly any difference between the ERPs regardless of whether the sentence was going to be rated as NotAcceptable or Acceptable. Intriguingly however, anomalous sentences with physical experiencer verbs would engender slightly more negative ERPs at the beginning of the experiment if the sentence was going to be rated as Acceptable as opposed to NotAcceptable, with this difference slowly but completely disappearing after about 1/3^rd^ of the total number of experimental trials. That is, even though the brains of the participants detected the anomaly in both violation conditions and therefore engendered a negativity, the behavioural ratings were not always a reflection this.

**Fig. 3.**
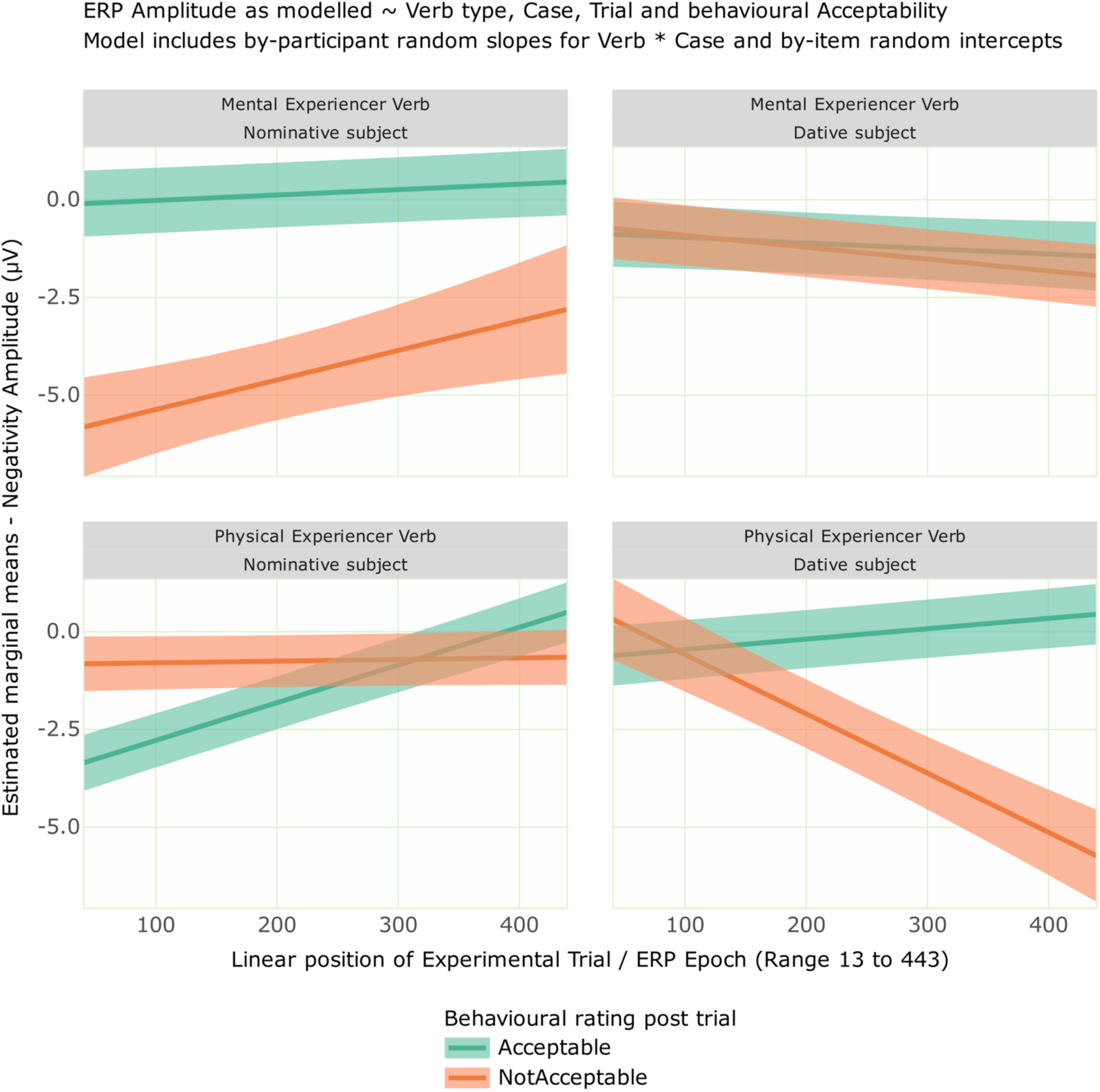
Estimated marginal means for the 400–800 ms negativity time-window by Verb type, Case, Trial and sentence-final behavioural Acceptability rating. Shaded areas represent 83% confidence intervals. Conditions plotted in the upper-left and lower-right quadrants are grammatical; conditions plotted in the upper-right and lower-left quadrants are ungrammatical.

More interestingly however, when a non-anomalous sentence of either verb type was going to be rated as NotAcceptable as opposed to Acceptable, a concomitant negativity ensued. This negativity was consistently large over the course of the experiment for mental experiencer verbs with (non-anomalous) nominative subjects, whereas it was almost non-existent at the beginning of the experiment for the physical experiencer verbs with (non-anomalous) dative subjects and grew larger and larger over the course of the experiment.

Simply put, the ERPs at the position of the experiencer verb are starkly different contingent upon the sentence-final acceptability for mental versus physical experiencer verbs. Surprisingly, this difference is observed for the non-anomalous conditions rather than for the anomalous conditions.

In order to examine whether this pattern is in some way a consequence of how the acceptability ratings for the critical conditions were distributed over the course of the experiment, Fig. 4 illustrates the trial-wise distribution of the acceptability ratings for the critical conditions. As can be inferred from Fig. 4, the seemingly ‘contradictory’ ratings (i.e., Acceptable ratings for ungrammatical sentences; NotAcceptable ratings for grammatical sentences) are spread over the entire course of the experiment more or less evenly rather than being clustered at a certain point in time during the experiment. So, this rules out a potential explanation for the pattern of results illustrated in Fig. 3 based on possible clustering differences of ratings for the different conditions over the course of the experiment.

**Fig. 4.**
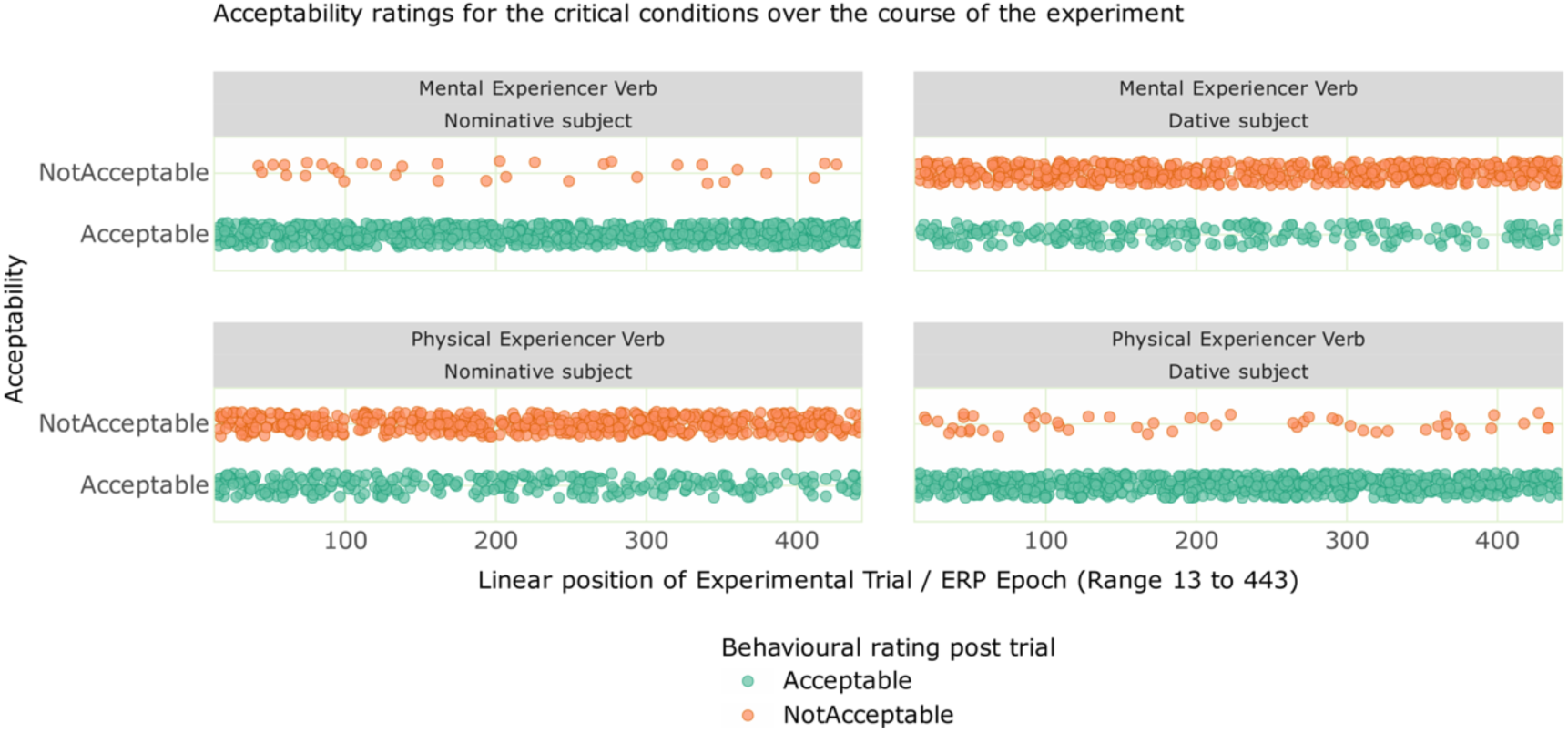
Trial-wise distribution of the acceptability ratings for critical sentences over the course of the experiment. Conditions plotted in the upper-left and lower-right quadrants are grammatical; conditions plotted in the upper-right and lower-left quadrants are ungrammatical.

A tabulation of the total number of trials that entered the analysis in each condition and the percentage of those trials that were rated as either Acceptable or NotAcceptable is shown in Table 3. This data shows that a relatively small number of trials from the non-anomalous conditions were rated as NotAcceptable, whereas a considerably larger number of trials with violation sentences were rated as Acceptable. Therefore, the ERPs for the violation conditions are not very different contingent upon sentence-final acceptability, in spite of a relatively higher number of such trials that were rated as Acceptable as opposed to NotAcceptable. That is, even though there may have been enough power in the signal for violation condition trials with Acceptable ratings compared to that for violation condition trials with NotAcceptable ratings, this did not however result in a concomitant difference in the ERPs for violation conditions contingent upon sentence-final acceptability ratings. In other words, all violations resulted in very similar ERPs (negativity) regardless of whether they were rated as Acceptable or NotAcceptable. By contrast, despite the fact that relatively few non-anomalous trials with NotAcceptable ratings are evenly and similarly distributed over the entire course of the experiment for both verb types (upper-left and lower-right quadrants in Fig. 4), they still resulted in very different ERP patterns for the two verb types (upper-left and lower-right quadrants in Fig. 3).

**Table 3.** Number and percentage of trials rated as Acceptable versus NotAcceptable in each critical condition.

| Case | Verb type | Total No. of Trials | No. of trials rated as Acceptable | No. of trials rated as NotAcceptable |
| --- | --- | --- | --- | --- |
| Nom | Mental | 793 | 759 (95.7 %) | 34 (4.3%) |
| *Dat | Mental | 799 | 238 (29.8 %) | 561 (70.2 %) |
| *Nom | Physical | 835 | 304 (36.4 %) | 531 (63.6 %) |
| Dat | Physical | 801 | 748 (93.4 %) | 53 (6.6 %) |

Seen in light of these details, the pattern of results in Fig. 3 qualifies the ERP plots in an important way and is indeed crucial in gaining a more nuanced understanding of the processing of the two verb types. Specifically, this pattern shows that the averaged ERP plots, albeit illustrating clearly that violations correlate with a negativity effect for both verb types, do not however capture the subtle but robust difference between processing mental experiencer and physical experiencer verbs, as evidenced by very different manifestations of the negativity when non-anomalous sentences were nevertheless contradictorily judged as NotAcceptable as opposed to Acceptable. This is clearly illustrated by the plots in the upper-left and lower-right quadrants of Fig. 3. Further, there is no evidence in the data for this difference to result from how the trial-wise acceptability ratings are distributed over the course of the experiment, for as Fig. 4 shows, the distribution of Acceptable and NotAcceptable ratings are distributed very similarly for both conditions of a given type (i.e., on the one hand, the two violation conditions pattern together; on the other, the two non-anomalous conditions pattern together).

Consequently, seen in conjunction with:

a. the different distribution of by-item acceptability ratings for the two verb types,
b. the similar trial-wise distributions of acceptability ratings for violations versus grammatical conditions, and
c. the differences found between the violation conditions on the one hand and the non-anomalous conditions on the other as regards how the ERPs for these conditions are contingent versus not contingent upon sentence-final acceptability ratings respectively, the stark difference apparent from Fig. 3 between mental experiencer verbs with nominative subjects versus physical experiencer verbs with dative subjects is suggestive of inherent qualitative differences in how the two types of experiencer verbs are processed in Malayalam.

### Summary of results

One, both violation conditions elicited a negativity effect between 400 and 800 ms after the onset of the verb.

Two, post-hoc analyses to examine the slightly different morphologies of the negativity effect between the verb types showed that the negativity effect for violations with mental experiencer verbs reached an earlier maximum, whereas the negativity effect for violations with physical experiencer verbs peaked later.

Three, an acceptability-contingent analysis of the ERPs showed that the sentence-final acceptability of trials modulated the ERPs for the non-anomalous conditions but not for the violation conditions. Further, this modulation was qualitatively different between mental and physical experiencer verbs. This difference ensued despite the very similar extent and distribution of acceptability ratings for these conditions over the course of the experiment.

## 6. Discussion

The present study examined how mental and physical subject experiencer verbs are processed in simple intransitive constructions in Malayalam. The behavioural task showed a direct relationship between grammaticality and acceptability, with violations leading to lower acceptability compared to their grammatically correct counterparts. This pattern was consistent for both mental and physical subject experiencer verbs, despite numerical differences in the mean acceptability between the violation conditions. Electrophysiological results at the verb for violation conditions of both verb types revealed a negativity effect in the 400–800 ms time window. The post-hoc analysis carried out by splitting this time window into two confirmed that the negativity effect reached an early maximum for mental experiencer verbs (400–600 ms), whereas it reached a later maximum for physical experiencer verbs (600–800 ms). The acceptability-contingent analysis revealed that the sentence-final acceptability of trials modulated the ERPs in non-anomalous conditions but not in violation conditions. Furthermore, it showed that this modulation qualitatively differed between mental and physical experiencer verbs. We however did not observe the thematic reanalysis related positivity effect that we predicted for the non-anomalous condition with mental experiencer verbs following a nominative subject. A more general language-internal motivation for the absence of positivity effects in our data is provided in the supplementary material, but the absence of a reanalysis related positivity is not surprising in view of the theoretical account of experiencer verbs in Malayalam mentioned earlier (Krishnan, 2019). According to this account, mental experiencer verbs pattern together syntactically with action verbs (i.e., they both require nominative subjects), because the subjects of these verb types have more control/agency over the action/state. Therefore, mental experiencer verbs, which involve a certain degree of agency compared to physical experiencer verbs, may not require thematic reanalysis. This could be attributed to the volitional nature of mental experiencer verbs, whereby the experiencer is (able to) willfully engage with (or choose not to engage with) experiencing the mental state expressed by the verb. Further, although Fig. 2 visually appears to suggest that the negativity for the physical experiencer verb violation condition was larger in amplitude compared to that for the mental experiencer verb violation condition, a closer examination of the statistical results in the analysis time window (400–800 ms) reveals that the mean amplitude for both violation conditions is, in fact, numerically almost equal: the estimate for the condition with mental experiencer verbs and dative subjects was -1.243 (SE = 0.569, p = 0.02, s = 5.11), whereas for the condition with physical experiencer verbs and nominative subjects, it was -1.126 (SE = 0.490 p = 0.02, s = 5.53). This overall pattern, i.e., negativities that are similar in amplitude for both violations, remains intact even when qualified by relative differences between the conditions in the split time-windows, in which we observed the negativity peak occurring earlier for mental experiencer verbs and later for physical experiencer verbs. Thus, in the earlier time-window in which the negativity peaked for the condition with mental experiencer verbs and dative subjects, the estimate for that condition was -1.235 (SE = 0.435, p = 0.004, s = 7.77); in the later time-window in which the negativity peaked for the condition with physical experiencer verbs and nominative subjects, the estimate for that condition was -1.337 (SE = 0.495, p = 0.006, s = 7.16). That is, the mean amplitude of the negativity was numerically similar for both verb types, with minimal difference between the two at their respective peaks.

Our results therefore show that both mental and physical experiencer verbs in Malayalam are processed qualitatively similarly, with violations of both verb types generally evoking a negativity effect with similar latency, amplitude and topography. Nevertheless, there are subtle differences between the verb types in terms of peak latency of the effect, i.e., when the negativity effect reaches its maximum peak, as well as in the acceptability contingent analysis. In what follows, we discuss the interpretations of the negativity effect that we found for both the experiencer verbs.

In the processing literature, negativity effects that are instances of the N400 component have been interpreted in various ways. Delogu et al., (2019) discuss three key theories about the functional interpretation of the N400 component. The access/retrieval account suggests that N400 component reflects the effort to access a word’s conceptual meaning from long-term memory, with more contextual cues reducing this effort (e.g., Brouwer et al., 2017; Kutas & Federmeier, 2011). The integration account posits that N400 component indicates the difficulty of integrating a word’s meaning into the existing context (e.g., Brown & Hagoort, 2000). The hybrid account combines these views, proposing that N400 reflects both retrieval and integration efforts (e.g., Baggio, 2018; Nieuwland et al., 2020). Bornkessel-Schlesewsky and Schlesewsky (2019) provide an interpretation of all language-related negativities based on the predictive coding framework, whereby the brain continuously generates predictions about upcoming linguistic input. The amplitude modulations of negativities including the N400 are said to reflect the degree to which these predictions are confirmed or violated. Importantly, the predictions are said to be precision-weighted, with precision reflecting the relevance of a particular stimulus feature in a given language. Previous studies on experiencer verbs have reported various ERP effects at the position of the verb, including N400, P600, early positivity, and LAN. Of relevance here, the N400 effect found in the context of experiencer verbs has been mainly attributed to grammatical function reanalysis based on the preferential ordering associated with such verbs (Bornkessel, et al., 2004; Schelesewsky & Bornkessel, 2006; Bornkessel et al., 2020), thematic-aspectual differences associated with them (Bourguignon et al. 2011), and violations of the verb’s selection restriction (Paczynski and Kuperberg, 2011).

The negativity observed in our study could be interpreted as resulting from an integration problem, when the parser encounters a mental or physical experiencer verb following a dative or nominative subject, respectively. This occurs because the lexical-semantic information of the verb cannot be easily integrated with the experiencer subject, as the two experiencer verbs constrain the case marking of the experiencer subject argument in specific ways (Friederici & Frisch, 2000; Horiguchi, 2019). More generally, our result could also be interpreted in terms of precision-weighted prediction errors occurring at the verb due to the mismatch between top-down expectations about the verb and the bottom-up information from the verb actually encountered (Bornkessel-Schlesewsky & Schlesewsky, 2019). The prominence information of the subject noun generates predictions about the lexical-semantic features of the upcoming verb, and when these predictions are not met, it leads to processing difficulties. Evidence from previous studies that included manipulations of morphological case, word order and verb type shows that, when there is a mismatch between the previously computed prominence and the establishment of the lexical-semantic structure of the verb, this may result either in an early parietal positivity (Bornkessel et al., 2003), a late parietal positivity (Friederici & Mecklinger, 1996; beim Graben et al., 2000), or in a centro-parietal negativity (Bornkessel et al., 2004; Leuckefeld, 2005). In our study, both violation conditions featured sentence-final verbs that contradicted the lexical-semantic expectations about the verb and resulted in a negativity effect.

A more pertinent interpretation for our result is along the lines of Choudhary et al. (2009) and Nieuwland et al. (2013), in which it was posited that violations of interpretively relevant linguistic rules result in an N400 effect. In Malayalam, a clear morphosyntactic constraint governs the case marking of subject experiencers in simple experiencer constructions based on the type of experience they convey. Specifically, subject experiencer verbs denoting mental experiences require nominative subjects, while those expressing physical experiences consistently require subjects marked with the dative case (Jayaseelan, 2004). This rule is interpretively relevant because the choice of case directly influences the interpretation of the verb’s meaning and reflects the structural distinction in the language. This interpretive relevance of the rule holds true equally for both verb types, as evidenced by very similar amplitudes of the negativity effect for both violation conditions.

A mental experiencer verb describes a state of change within an individual, and the individual (i.e., the experiencer argument) has more control over the state such that they can volitionally effect a change or influence it. For example, it is possible for the experiencer to willfully distract themselves from the happiness that they are experiencing, say, in order to become more balanced in their outlook or to suppress expressions of their happiness in front of others. That is, the experiencer argument has volitional control to effect a change in their mental experience without taking any tangible action. This potential for volitional control is marked by the nominative case of the mental experiencer argument. On the other hand, a physical experiencer verb describes the advent of a state, and the experiencer has hardly any control over the state and cannot effect a change in the state of affairs per se, without taking further tangible action specifically towards mitigating the physical experience. For example, a person experiencing extreme cold in freezing temperatures cannot, under normal circumstances, distract themselves to feel less cold, unless they take some tangible action (such as going indoors, wearing warm clothes etc.) towards achieving this goal. More compelling examples of this sort would be the physical experience of pain or hunger. The lack of volitional control over the state of affairs in these examples requires marking the experiencer argument with the dative case in Malayalam (Mohanan and Mohanan, 1990, Jayaseelan, 2004; Krishnan, 2019). In other words, marking a subject argument nominative versus dative in Malayalam is not arbitrary; it is interpretively highly relevant, and is intended to directly convey information about the potential availability of volitional control for the experiencer argument versus lack thereof.

As per this rule, the nominative and dative subjects in our study would have given rise to different expectations about the upcoming verb. Nominative subjects would be compatible with a range of verb types, whereas dative subjects would only be compatible with a restricted set of verb types that specifically indicate a lack of volitional control on the part of the subject argument. When the parser encounters a physical experiencer verb following a nominative subject, or a mental experiencer verb following a dative subject, as in the violation conditions in our study, the expectations generated by the subject case about the verb are not met. That is, these constitute a violation of the interpretively relevant rule that specifies the subject case based on whether or not the verb entails volitional control of the subject, which then results in engendering a negativity effect in comparison to the non-anomalous conditions. We interpret this negativity as an instance of the N400 component, in line with similar interpretations from other typologically unrelated languages such as Hindi (Choudhary et al, 2009) and Spanish (Nieuwland et al., 2013). Crucially, there was no evidence for amplitude modulations of the negativity between the two verb types in our data. This reflects the fact that the subject marking rule is equally relevant for both mental as well as physical experiencer verbs. The negativity effect found in our study showed a relatively long latency for both mental and physical experiencer verbs. Indeed, such long-latency negativities have previously been reported in several studies, both in the auditory as well as visual modalities, due to word variability across trials (Holcomb and Neville, 1990), post-lexical integration of words into higher order meaning (e.g., Brown and Hagoort, 1993; Holcomb, 1993; Steinhauer et al., 2017), complex processing required for mismatches involving verbs, which may be more demanding than those involving nouns (Courteau et al., 2019), unagreement (Ito et al., 2020), agreement violations (Bhattamishra et al., 2020), and reversal anomalies (Bornkessel-Schlesewsky et al., 2011).

Despite the fact that violations of both verb types elicited a negativity effect, there are nevertheless subtle but robust differences between the processing of the two verb types that came to light in our post-hoc analysis. A first such difference is the peak latency of the negativity effect, which occurred earlier for the mental experiencer violations but later for the physical experiencer violations. Note that this difference could be observed only when the original analysis time-window spanning from 400 ms to 800 ms after the onset of the verb was split into two halves, and that the overall effect is present for both verb types in the full time-window of analysis. This is in line with our hypothesis that processing violations involving physical experiencer verbs would be more costly than those with mental experiencer verbs, although we predicted that this would lead to amplitude differences (which is not the case; see above) rather than to a peak latency difference between the verb types. We speculate that a general difference in the possible range of expectations that a nominative versus a dative subject generates could explain the observed ERP peak latency difference between the verb types. As mentioned earlier, nominative subjects are compatible with a vast range of verb types, and therefore the range of possible expectations that nominative subjects generate is very broad, and in turn less specific and precise. By contrast, dative subjects in Malayalam are only compatible with a restrictive set of stative constructions, amongst which physical experiencer verbs are much more likely, as cross-linguistic studies show that dative subjects are more commonly associated with experiencer verbs (Van Valin, 1991). Consequently, the range of expectations about the verb that a dative subject generates is much more specific and precise. Seen in light of this, for violations involving mental experiencer verbs following dative subjects, the verb disconfirms a very specific expectation for the verb type, and therefore a negativity ensues and peaks relatively early on. By contrast, for violations involving physical experiencer verbs following nominative subjects, the verb disconfirms a less specific expectation. In terms of the neurobiologically plausible model of language-related negativities proposed by Bornkessel-Schlesewsky & Schlesewsky (2019), the fact that nominative case is the default subject case for action verbs, and thus for the vast majority of verbs, adds to the decreased precision with which it can predict a specific kind of verb. Further, given that the other type of experiencer verbs (namely mental experiencer verbs) is indeed compatible with nominative subjects makes the anomaly more complex. This could potentially account for the later maximum of the negativity effect for physical experiencer verbs with a nominative subject.

This tentative explanation notwithstanding, more work is necessary in this regard. A limitation of our study is that it is not possible to disentangle the effect of differing expectations due to nominative versus dative subjects interacting with the two types of experiencer verbs. One potential way to mitigate this limitation could involve examining mental and physical experiencer verbs within complex constructions, because the subject argument in such constructions require dative subjects for both mental and physical experiencer predicates (see examples 11 and 12). Such an investigation could help shed light on whether the observed differences in processing are driven exclusively by the verb type or not.

Nonetheless, converging evidence in support of the argument that the processing of the two verb types is indeed subtly different stems from the distinct pattern of results found for mental and physical experiencer verbs in the analysis of the ERPs contingent upon sentence acceptability. It is striking that it was the non-anomalous sentences rated as NotAcceptable that showed a clear difference between the verb types in this analysis. Indeed, the ERPs were consistently more negative for non-anomalous sentences with a mental experiencer verb following a nominative subject when they were rated as NotAcceptable as opposed to Acceptable. This difference remained prominent across trials throughout the course of the experiment. In stark contrast, the ERPs for non-anomalous sentences with a physical experiencer verb following a dative subject showed no such difference in the beginning of the experiment, regardless of whether these were rated as NotAcceptable or Acceptable. Subsequently however, a gradual difference emerged such that the ERPs were more negative for the non-anomalous condition with physical experiencers when these were rated as NotAcceptable as opposed to Acceptable. Crucially, as the bottom right panel in Fig. 3 shows, the magnitude of this difference steadily increased over the course of the experiment, becoming ever more prominent towards the end. As discussed earlier, there is no evidence in our data for an explanation for this asymmetric difference in the pattern of results between the two verb types based on potential clustering differences of ratings for the different conditions over the course of the experiment. This is because, as illustrated in Fig. 4, the contradictory ratings are spread evenly across the entire experiment rather than being clustered at certain points.

The finding is particularly intriguing because the aforementioned difference in the ERPs between mental and physical experiencer verbs in non-anomalous sentences emerged from a relatively small number of trials that were rated as NotAcceptable. Furthermore, this pattern is in contrast to the ERPs for the violation conditions, which are not very different for the two verb types contingent upon sentence-final acceptability, in spite of a relatively higher number of such violation trials that were contradictorily rated as Acceptable as opposed to NotAcceptable. In short, neither the number of contradictory ratings, nor how they are spread over the course of the experiment could account for the differences observed. Instead, this pattern of results is indicative of subtle differences inherent in the processing of non-anomalous sentences of the two verb types.

## 7. Conclusion

The present study explored the processing of mental and physical experiencer verbs in Malayalam. The experiment revealed a negativity for subject case violations involving both mental and physical experiencer verbs, suggesting that different experiencer verb types are processed qualitatively similarly. However, post-hoc analyses of the data revealed more nuanced ERP differences between the two verb types. Specifying a more precise mechanism underlying the observed differences is outstanding, and further research is necessary to shed light on this aspect. Nevertheless, despite the fact that both mental experiencer verbs and physical experiencer verbs are processed qualitatively broadly similarly, there is sufficient evidence in our data to conclude that subtle but robust differences are inherent in processing mental versus physical experiencer verbs in Malayalam.

## Data availability

Data repository. The raw data collected during the experiment, the pre-processing pipeline used and the pre-processed data are available here: < https://doi.org/10.5281/ZENODO.14986232>

Analysis repository. The analysis code and full model outputs of all the analyses reported are available as R notebooks here: <Zenodo.org link>

(Data and analysis code will be made available on Zenodo.org, after the final acceptance of the article.)

## Appendix / Supplementary Material

### A1. ERPs at the sentence-initial subject noun

Fig. S1 shows the ERPs at the position of the sentence-initial subject noun collapsed over verb type, since the verb is yet to unfold following the noun. A linear mixed effects model similar to model m1 (see the planned analysis of the verb in the main article) with maximal random effects specification for the single-trial ERP amplitudes at the position of the noun was computed in three time-windows, namely 150–350 ms, 400–600 ms and 600–800 ms. The analysis code and full model outputs are available as R notebooks in the analysis repository online. Type II Wald Chi-squared tests on this model in the respective time-windows showed that there were no main effects nor interactions involving the factor Verb type, which is unsurprising given that the verb is yet to unfold in the sentence following the noun. This result supports visualising the ERPs at the noun collapsing over Verb type, as shown in Fig. S1. There was an interaction of ROI x Case in the 150–350 ms and 400–600 ms time-windows, showing that the physical (orthographic) difference between the dative and nominative subjects was perceived (in the 150–350 ms time-window, typical for P200 effects) and that the two types of subjects evoked different effects at the lexical processing stage (in the 400–600 ms time-window). There were no effects in the 600–800 ms time-window. Whilst we refrain from interpreting these findings further in view of the fact that they did not directly form part of our hypotheses, they nevertheless are suggestive of processing differences between dative subjects and nominative subjects both at the pre-lexical as well as post-lexical stages, which is in general in line with findings from Tamil, a related Dravidian language (Muralikrishnan, 2024), albeit with crucial differences perhaps due to modality differences between the Tamil study and our study. If dative and nominative subjects are processed differently, then it is indeed plausible that they also give rise to different expectations about the upcoming verb.

**Fig. S1.**
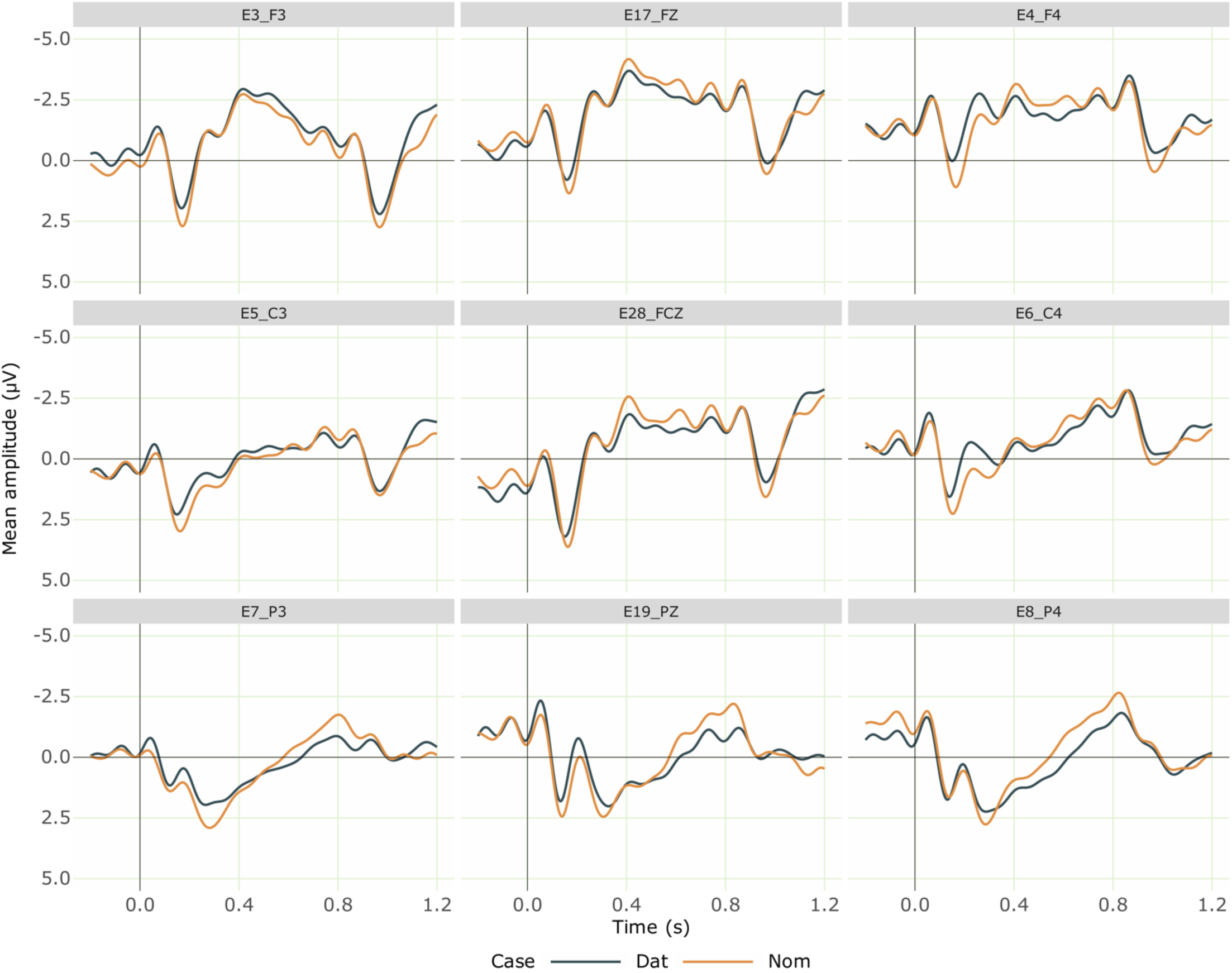
Grand averaged ERPs at the nominative and dative sentence-initial subject nouns from 28 participants (collapsed over verb type, since the verb is yet to unfold following the noun). Negativity is plotted upwards; the time axis runs from -0.2 s to 1.2 s (i.e., -200 ms to 1200 ms) with 0 being the onset of the noun. The dark blue line shows the ERPs for the nominative nouns and the orange line shows that for the dative nouns.

### A2. Absence of positivities at the Verb

Fig. S2 and Fig. S3 show the ERPs at the verb for mental and physical experiencer verb conditions separately respectively. We did not observe late positivity effects in violation conditions involving neither mental nor physical experiencer verbs. This is despite the fact that ill-formed sentences typically tend to elicit a P600 effect (Bornkessel-Schlesewsky & Schlesewsky, 2008), and in spite of the presence of a judgement task that is known to modulate the amplitude of late positivities ( Bornkessel-Schlesewsky, 2011). We speculate that this is due to the resolvability of the violation conditions in our study independent of conflict strength (Frenzel et al., 2011). The combination of nominative subjects and physical experiencer verbs can be non-anomalous in Malayalam when the sentence fragment continues further to express reported speech or a figurative meaning. That is, several non-anomalous continuations are possible for the physical experiencer violation condition in our study. This is illustrated by the example sentences A and B, which are well-formed and felicitous sentences in Malayalam. For the combination of dative subjects and mental experiencer verbs, such continuations are not possible without orthographic changes in the verb form. Nevertheless, there is an emerging phenomenon in everyday language use, which may be relevant here. The enunciative word final schwa /ә/ in Malayalam (or its variants /u/ and /ɯ/ in other Dravidian languages), commonly employed in to phonologically adapt non-Dravidian words to conform with Dravidian phonological rules, has been observed to be regularly dropped under certain circumstances (Namboodiripad, 2021). A sort of an opposite of this phenomenon is also observable in native adverbial participles, which are very frequent in Malayalam. To derive a non-finite adverbial participle from a finite verb, if the final vowel in the past tense form of the finite verb is /u/, then this has to be changed to /ә/ to express the adverbial meaning. Crucially, the /ә/ in these verb forms are not enunciative, but morphemic, and obligatory as per the prescriptive grammar. However, in everyday use in orthography (but, importantly, not in speech!), this /ә/ often gets replaced with /u/. In essence, the adverbial form is represented as the finite past tense form due to the orthographic change. Anecdotal evidence suggests that this ‘alternative’ is quite frequent, and readers do not perceive the difference unless specifically asked about it. This has consequences for our stimuli, because the verbs in our stimuli were all in the past tense, and all of these past tense forms ended in /u/. If the orthographic change described above is so common as the anecdotal evidence suggests, especially with young readers, then it is possible that the verbs in our study (including the mental experiencer verbs) could potentially be read as the ‘alternative’ adverbial form, and then sentence continuations are always possible. Simply put, this emerging orthographic variation in everyday use could potentially create the illusion that the sentences could continue as constructions with adverbial verbs, with the final verb yet to occur. In case of our mental experiencer condition with dative subjects, these continuations could be, for instance, modal constructions such as the example sentences in C and D (a # indicates the ‘alternative’ form). Whilst this is a phenomenon that is not yet fully grammaticalized, this could nevertheless potentially explain why sentence continuations, and therefore possibilities for alternative interpretations, are still open on encountering the violating verb.

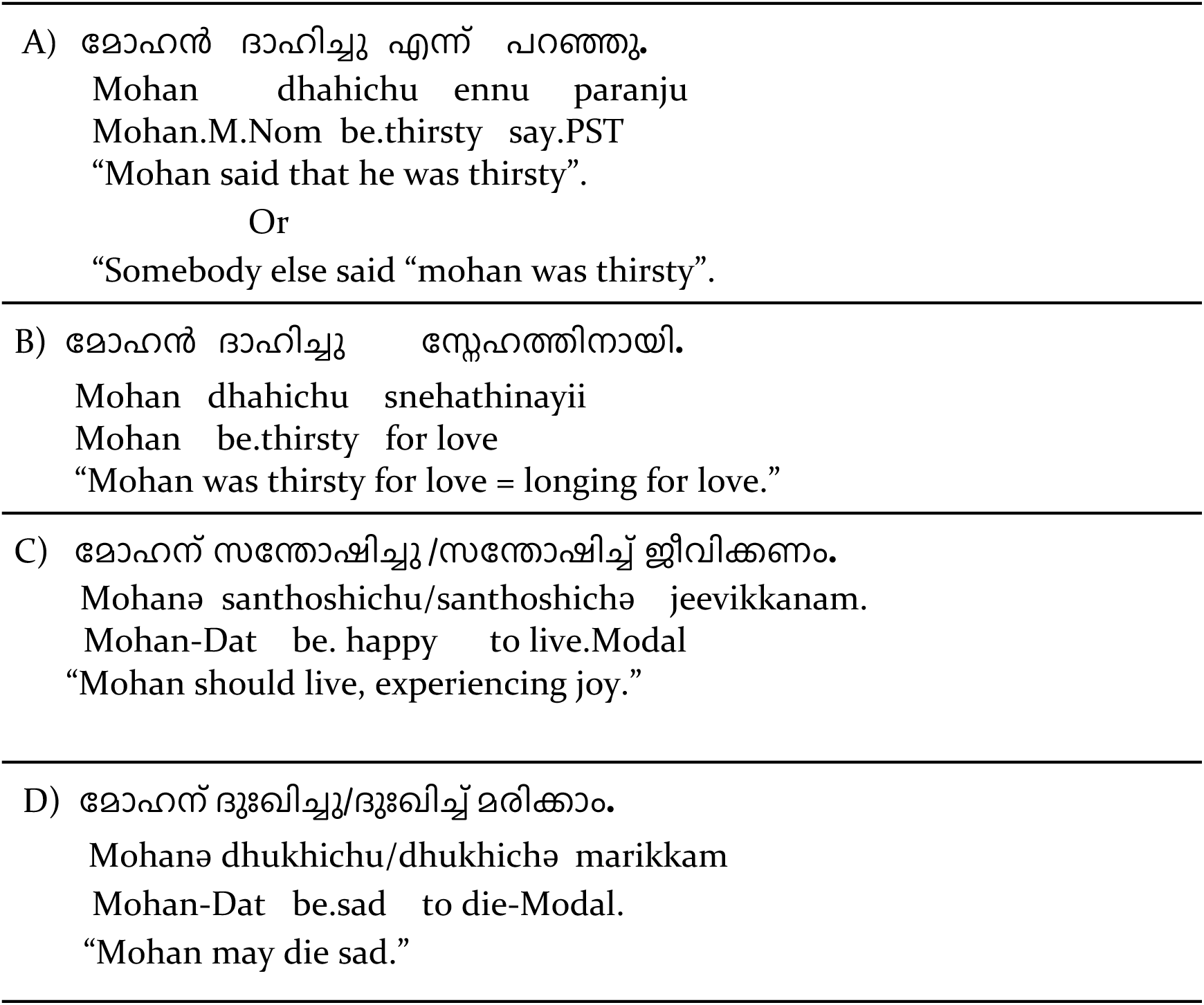

**Fig. S2.**
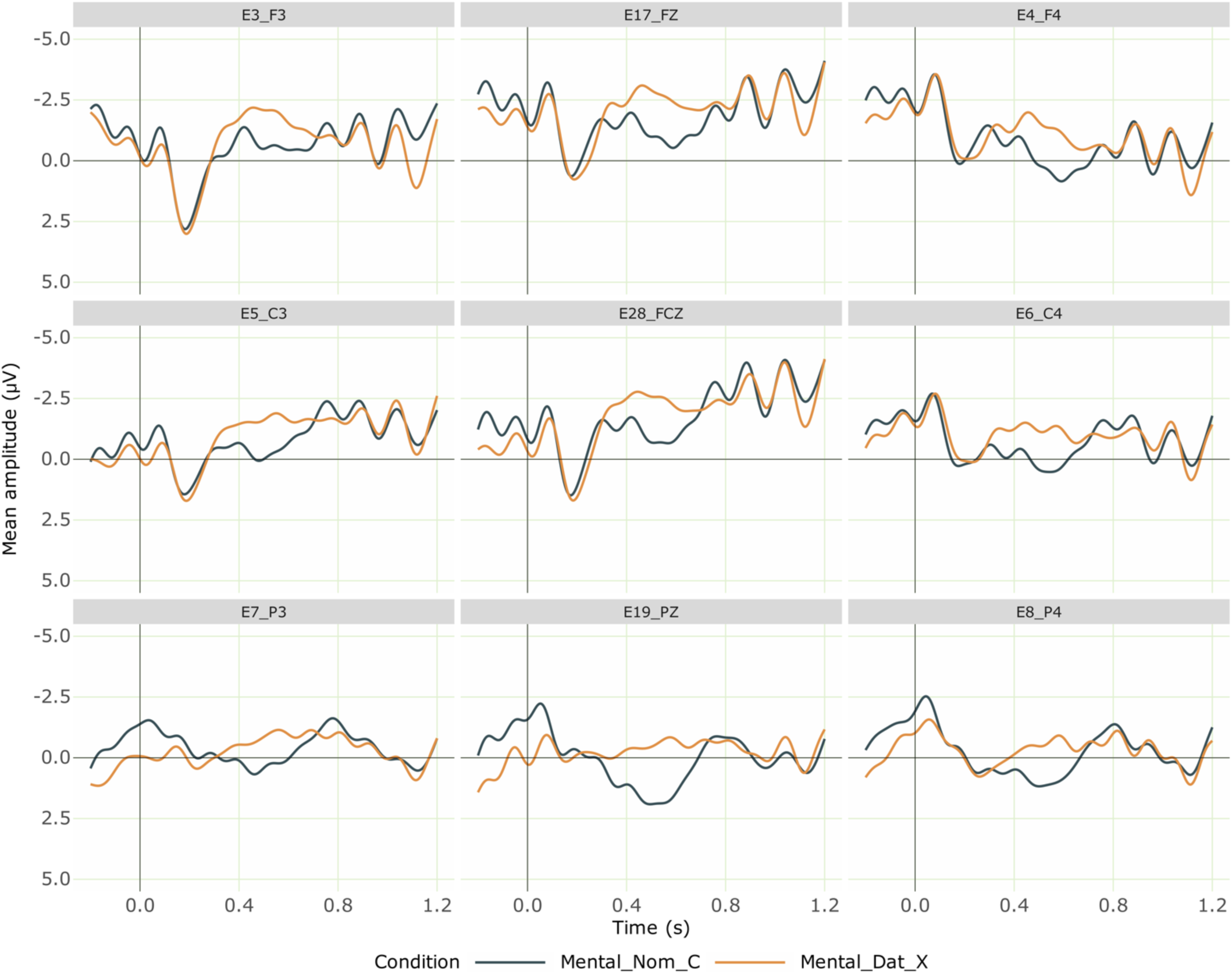
Grand averaged ERPs at the position of Mental experiencer verb comparing Nominative mental experiencer verb (Correct) and Dative mental experiencer verb (Incorrect) from 28 participants. Negativity is plotted upwards; the time axis runs from -0.2 s to 1.2 s (i.e., -200 ms to 1200 ms) with 0 being the onset of the critical verb. The dark blue line shows the ERPs for the correct Mental experiencer verb (with a nominative subject) and the orange line shows that for incorrect mental experiencer verb (with a dative subject), which elicited a negativity effect.

#### Mental experiencer verb

#### Physical experiencer verb

**Fig. S3.**
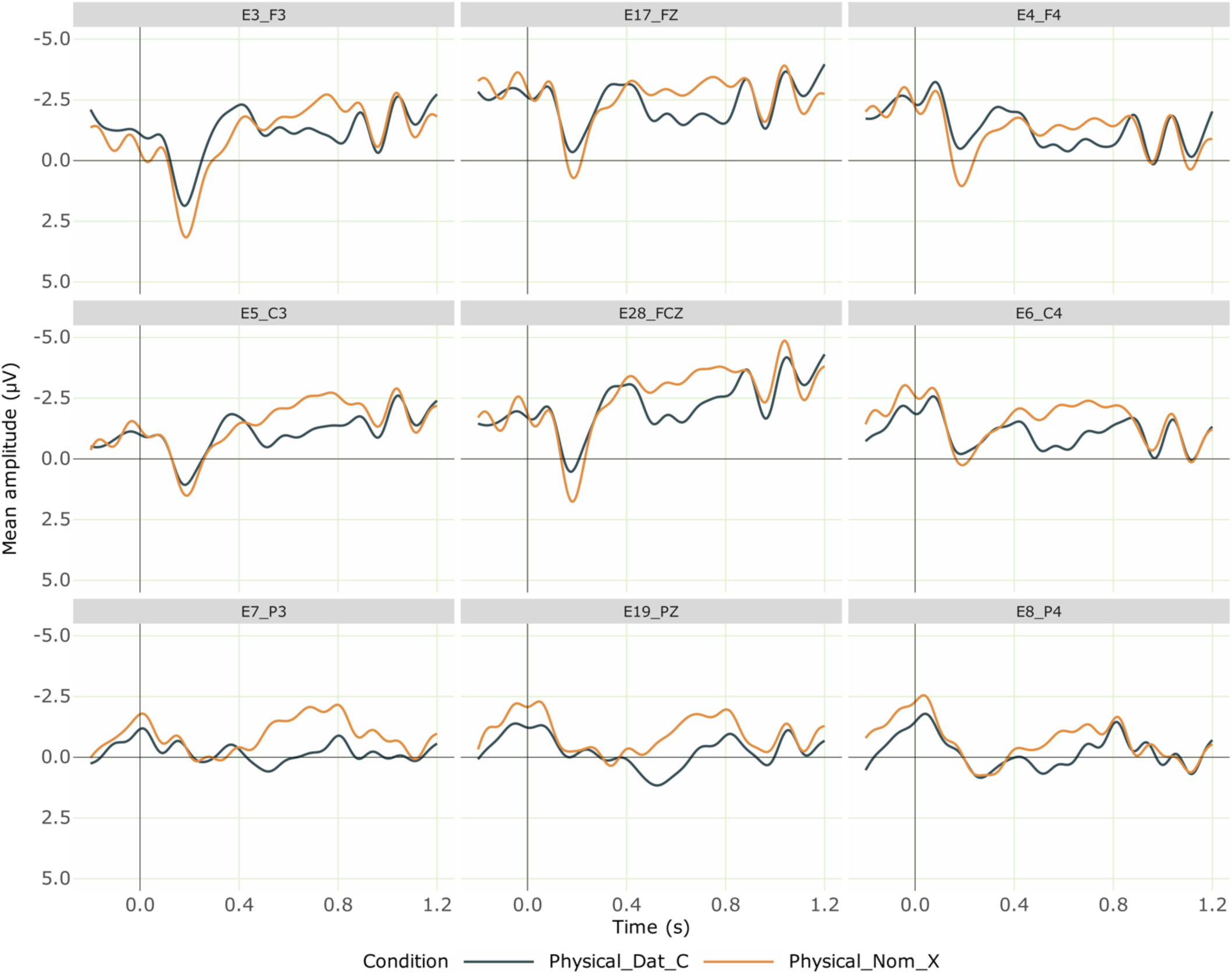
Grand averaged ERPs at the position of Physical experiencer verb comparing Dative Physical experiencer verb (Correct) and Dative Physical experiencer verb (Incorrect) from 28 participants. Negativity is plotted upwards; the time axis runs from -0.2 s to 1.2 s (i.e., -200 ms to 1200 ms) with 0 being the onset of the critical verb. The dark blue line shows the ERPs for the correct Physical experiencer verb (with a dative subject) and the orange line shows that for incorrect Physical experiencer verb (with a nominative subject), which elicited a negativity.

## Footnotes

1 The usage of these case suffixes is based purely on the phonology (last syllable) of the noun to which these are attached, regardless of the gender, number and other properties of the noun concerned. The allomorph “-ə” is used for nouns or their derivations ending in “n”, while “-kkə” is applied elsewhere (Rajendran, 1977). For example, for a noun ending in “-n” in singular (such as nadan – actor), the appropriate dative case marker would be “-ə”, thus yielding “nadanə” – actor.DAT. But when the same noun is in plural, as in “nadanmaar – actors”, then the correct dative case marker would be “-kkə, thus yielding “nadanmaarkkə - actors.DAT”, because the plural noun form does not end in “-n”. Therefore, the choice of the allomorph is purely based on the final phoneme in a noun or its derived form to which the dative case marker morpheme should be suffixed.

